# Elamipretide Improves ADP Sensitivity in Aged Mitochondria by Increasing Uptake through the Adenine Nucleotide Translocator (ANT)

**DOI:** 10.1101/2023.02.01.525989

**Authors:** Gavin Pharaoh, Varun Kamat, Sricharan Kannan, Rudolph S. Stuppard, Jeremy Whitson, Miguel Martin-Perez, Wei-Jun Qian, Michael J. MacCoss, Judit Villen, Peter Rabinovitch, Matthew D. Campbell, Ian R. Sweet, David J. Marcinek

## Abstract

Aging muscle experiences functional decline in part mediated by impaired mitochondrial ADP sensitivity. Elamipretide (ELAM) rapidly improves physiological and mitochondrial function in aging and binds directly to the mitochondrial ADP transporter ANT. We hypothesized that ELAM improves ADP sensitivity in aging leading to rescued physiological function. We measured the response to ADP stimulation in young and old muscle mitochondria with ELAM treatment, *in vivo* heart and muscle function, and compared protein abundance, phosphorylation, and S-glutathionylation of ADP/ATP pathway proteins. ELAM treatment increased ADP sensitivity in old muscle mitochondria by increasing uptake of ADP through the ANT and rescued muscle force and heart systolic function. Protein abundance in the ADP/ATP transport and synthesis pathway was unchanged, but ELAM treatment decreased protein s-glutathionylation incuding of ANT. Mitochondrial ADP sensitivity is rapidly modifiable. This research supports the hypothesis that ELAM improves ANT function in aging and links mitochondrial ADP sensitivity to physiological function.

Graphical Abstract.
ELAM Binds Directly to ANT and ATP Synthase and ELAM Treatment Improves ADP Sensitivity, Increases ATP Production, and Improves Physiological Function in Old Muscles.
ADP (adenosine diphosphate), ATP (adenosine triphosphate), VDAC (voltage-dependent anion channel), ANT (Adenine nucleotide translocator), H^+^ (proton), ROS (reactive oxygen species), NADH (nicotinamide adenine dinucleotide), FADH_2_ (flavin adenine dinucleotide), O_2_ (oxygen), ELAM (elamipretide), −SH (free thiol), −SSG (glutathionylated protein).

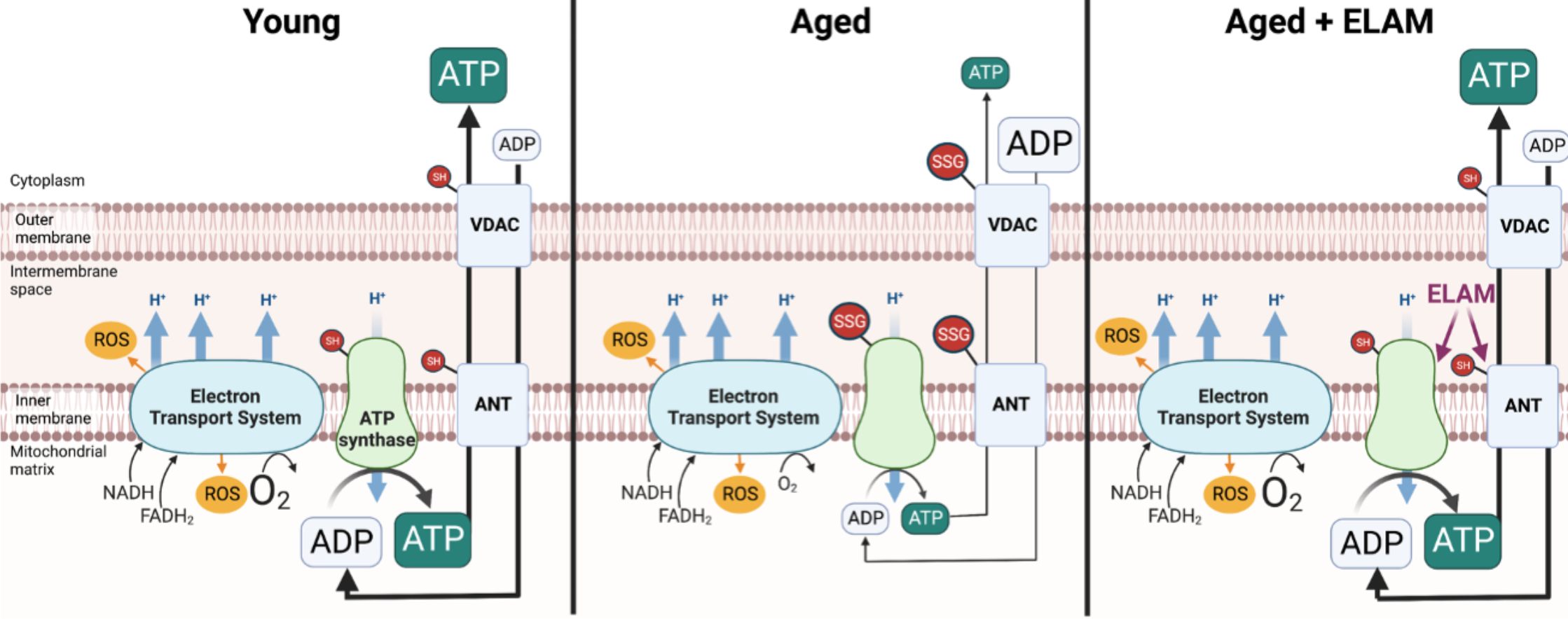

## INTRODUCTION

Mitochondrial dysfunction is a major hallmark of aging and has particular relevance in muscle aging (Lopez-Otin et al., 2013; Peterson et al., 2012). Declines in mitochondrial function are a primary mediator of age-related loss of muscle mass and strength (Peterson et al., 2012). Many parameters of mitochondrial function change with age and have been extensively reviewed (Sun et al., 2016). Loss of adenosine diphosphate (ADP) sensitivity, otherwise known as ADP insensitivity, is a key emerging phenotype of aging muscle (Chavez et al., 2020; Gouspillou et al., 2014; Holloway et al., 2018; Pharaoh et al., 2021). ADP insensitivity, assessed by titrating ADP into samples and measuring mitochondrial responses, is when a given concentration of ADP stimulates a smaller change in mitochondrial metabolism. ADP insensitivity impairs mitochondrial function in aging muscle including reports of decreased OCR and ATP production and increased reactive oxygen species (ROS) production at physiologically relevant concentrations of ADP (Chavez et al., 2020; Gouspillou et al., 2014; Holloway et al., 2018; Pharaoh et al., 2021). ADP is transported into the mitochondrial matrix by the voltage-dependent anion channel (VDAC) in the outer mitochondrial membrane and the adenine nucleotide translocator (ANT) in the inner membrane (**Graphical Abstract**) (Holloway et al., 2018). Importantly, changes in VDAC or ANT protein content cannot adequately explain the changes observed in ADP sensitivity (Gouspillou et al., 2014; Holloway et al., 2018; Miotto et al., 2018; Pharaoh et al., 2021). ADP sensitivity can be rapidly modified by acute exercise or insulin treatment within one hour, which suggests that regulation is likely at the post-translational rather than transcriptional or translational levels, although the exact mechanism is still not understood (Barbeau et al., 2018; Miotto et al., 2018).

Elamipretide (ELAM; previously known as SS-31 or Bendavia) is a mitochondrial-targeted peptide that rapidly improves physiological function in aging and mitochondrial myopathies (Campbell et al., 2019; Chiao et al., 2020; Reid Thompson et al., 2021; Siegel et al., 2013; Whitson et al., 2020). ELAM exerts multifaceted effects on mitochondrial function through interactions with cardiolipin (CL) and cardiolipin-associated proteins (Allen et al., 2020; Chavez et al., 2020; Mitchell et al., 2020). Chronic ELAM administration (8 weeks to 48 weeks) increases exercise tolerance, muscle mass, fatigue resistance, and *in vivo* ATPmax (Campbell et al., 2019; Reid Thompson et al., 2021). Other studies have shown beneficial effects on cardiac diastolic function in aging mice (Chiao et al., 2020; Whitson et al., 2020). Surprisingly, acute administration (1 hr) still improves muscle fatigue resistance and improves muscle ATPmax and ADP sensitivity *in vivo* in aged mice (Chavez et al., 2020; Siegel et al., 2013). In aged mouse muscle resting ADP concentration (~5-100 μM) increases with age *in vivo*, and ELAM treatment partially restores ADP concentration to young levels (Campbell et al., 2019; Siegel et al., 2013). We recently identified that ELAM binds directly to key components of the ADP/ATP transport and production pathways including ANT and ATP Synthase as well as 10 other mitochondrial proteins including proteins involved in ATP transport, the electron transport system, and glutamate metabolism (Chavez et al., 2020). In addition to direct protein binding, ELAM treatment stabilizes the ATP Synthasome, a protein supercomplex including ANT, ATP synthase, creatine kinase, and inorganic phosphate carrier and reduces proton leak through the ANT in aged cardiomyocytes (Zhang et al., 2020). In an oxidative stress-induced mouse model of sarcopenia, ELAM treatment repaired the sensitivity to ADP (Pharaoh et al., 2021).

Together, these data suggested ELAM as a prime candidate to improve ADP sensitivity in aging. We hypothesized that ELAM improves ADP sensitivity by improving ANT function and increasing ADP uptake (**Graphical Abstract**). To test this hypothesis, we measured the response to ADP stimulation in young and old mitochondria with ELAM treatment for all components of the ADP/ATP transport and synthesis pathway including ADP uptake, respiration (OCR), ROS production, mitochondrial membrane potential, and ATP production. In addition, we measured *in vivo* heart and muscle function and compared protein abundance, phosphorylation, and oxidative modification of ADP/ATP pathway proteins with age and ELAM treatment.

## RESULTS

### Respiration and Membrane Potential Response to ADP Stimulation in Isolated Muscle Mitochondria Require Supplemental Cytochrome c and Hexokinase Clamp

A major goal of this study was to measure mitochondria functional responses as a function of ADP concentration. To do this, we first validated assay conditions. We measured the response to ADP titration in isolated muscle mitochondria across a range of physiological ADP concentrations for oxygen consumption rate (OCR), membrane potential, and ROS production rate. Almost all previous studies on ADP sensitivity used permeabilized muscle fibers as a model with a focus on OCR. Developing isolated mitochondria as a model for these experiments allows the simultaneous measurement of many variables from the same sample as well as the use of mitochondria from tissues other than skeletal muscle. We addressed two limitations in isolated muscle mitochondria as a model to measure ADP sensitivity. First, isolated mitochondria rapidly consume ADP at subsaturating concentrations, a phenomenon first identified in 1955 as the state 3/4 transition by Chance and Williams (Chance and Williams, 1955). To allow steady-state measurements at subsaturating ADP concentrations as low as 1 μM, we used the hexokinase clamp system to consume the ATP produced by the mitochondria to maintain ADP at a steady-state level (Lark et al., 2016) (**Figure 1A**). The addition of the hexokinase clamp to isolated muscle mitochondria greatly decreased the EC_50_ for OCR and IC_50_ for membrane potential (**Figure 1B-C, 1E-F**). The second limitation is mitochondrial isolation from skeletal muscle using differential centrifugation damages the outermembrane, causing cytochrome c release that inhibits respiration capacities. By adding exogenous cytochrome c before measurement of ADP titrations, we removed this limitation and measured the full respiration capacities (**Figure 1A**). Exogenous cytochrome c increased respiration capacity but did not affect maximum membrane potential (**Figure 1D, 1G**). Both hexokinase clamp and cytochrome c addition were required to measure kinetic and maximum responses to ADP stimulation in mitochondria, respectively.

**Figure 1.**
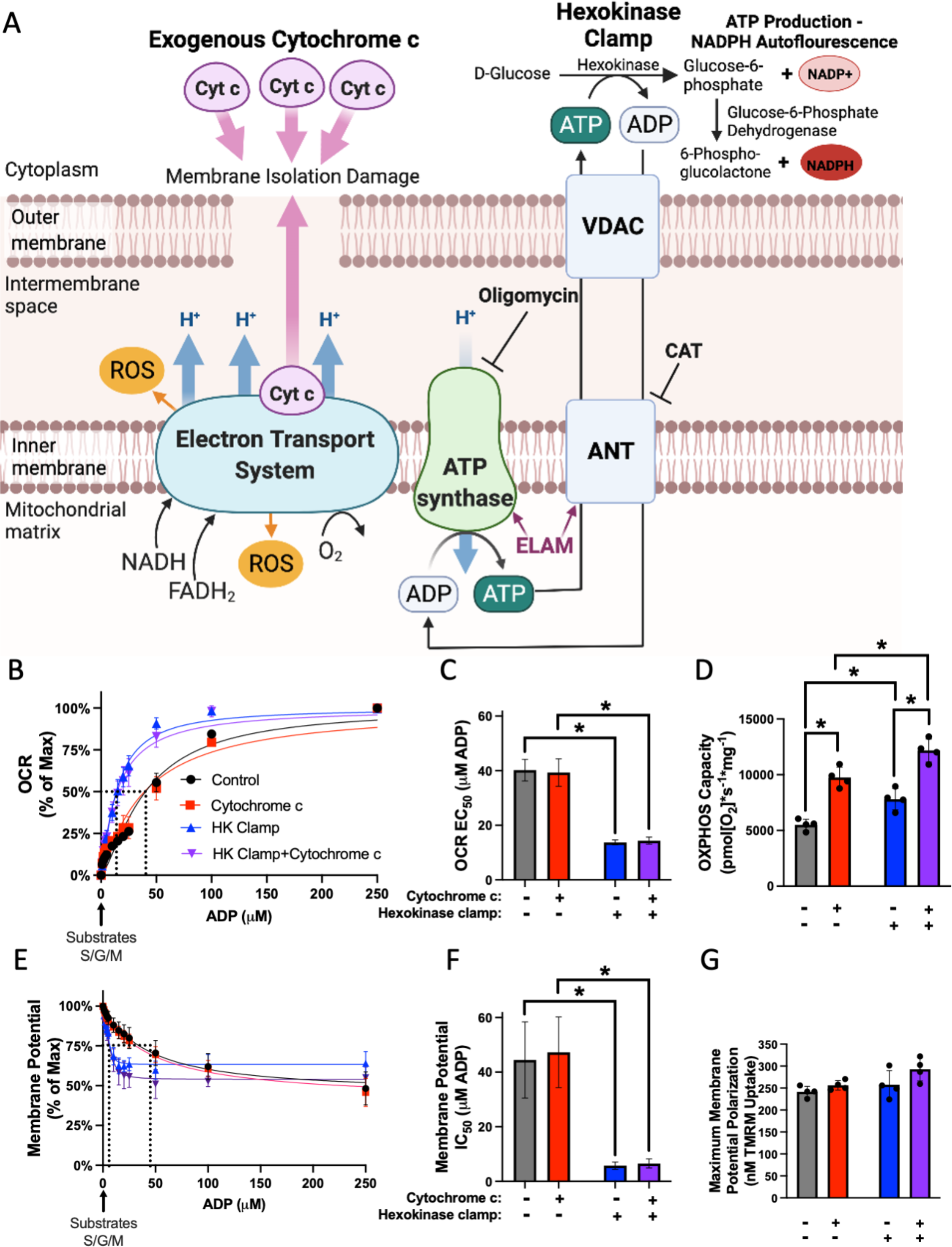
Respiration and Membrane Potential Response to ADP Stimulation in Isolated Muscle Mitochondria Require Supplemental Cytochrome c and Hexokinase Clamp. A) ADP/ATP transport pathway and binding of ELAM to ATP Synthase and ANT. Hexokinase clamp (1 U/mL hexokinase, 2.5 mM D-glucose) replenishes ADP for respiration at low ADP concentrations and exogenous cytochrome c (10 μM) prevents limitation on respiration from cytochrome c leak. ANT (Adenine nucleotide translocator), VDAC (Voltage-dependent anion channel), ROS (Reactive oxygen species), CAT (carboxyatractyloside). Generated with BioRender. B) OCR response curve to ADP, B) Calculated EC_50_, and (C) Maximum respiration capacity in isolated muscle mitochondria with control, 10 μM exogenous cytochrome c, hexokinase (HK) clamp, or both HK Clamp and cytochrome c conditions from young (5–8 mo) male C56Bl6/J mice (n=4). D) Membrane potential response curve to ADP, E) Calculated IC_50_, and F) Maximum membrane potential in isolated muscle mitochondria with control, 10 μM exogenous cytochrome c, hexokinase (HK) clamp, or both HK Clamp and cytochrome c conditions from young 5–8 mo male C56Bl6/J mice (n=4). Significance determined by Two-way RM ANOVA with Tukey’s posthoc test, *p<0.05. Mean ± SD.

### ELAM Increases Sensitivity to ADP for Respiration in Old Muscle Mitochondria

We dissected hindlimb muscles from young and old male mice and isolated mitochondria from one hindlimb using control buffers and the other hindlimb in control buffers supplemented with ELAM (acute treatment) (**Figure 2B**). We also treated old (26 mo to 28 mo) male mice with ELAM for 8 weeks by osmotic minipump, then isolated their hindlimb muscle mitochondria with ELAM (chronic treatment) (**Figure 2B**). We measured the response to different ADP concentrations by these mitochondria reflected by parameters of mitochondrial function, including respiration, membrane potential, ROS production, ATP production, and ADP uptake (**Figure 2A-B**). A unique aspect of this study is the detailed simultaneous characterization of mitochondrial samples from the same animal that allows for a comparison of how several variables of mitochondrial function directly relate. Using principal component analysis (PCA) of the multivariate mitochondrial data, aged mitochondria chronically treated with ELAM separated distintinctly from young and old mitochondria (**Figure S1A**). In the leak state, OCR was significantly correlated with mitochondrial membrane potential (p<0.05 by linear regression) (**Figure S1B**). While leak OCR was not significantly correlated with ROS production, leak ROS production and membrane potential were correlated (p<0.0001 by linear regression) (**Figure S1C-D**).

**Figure 2.**
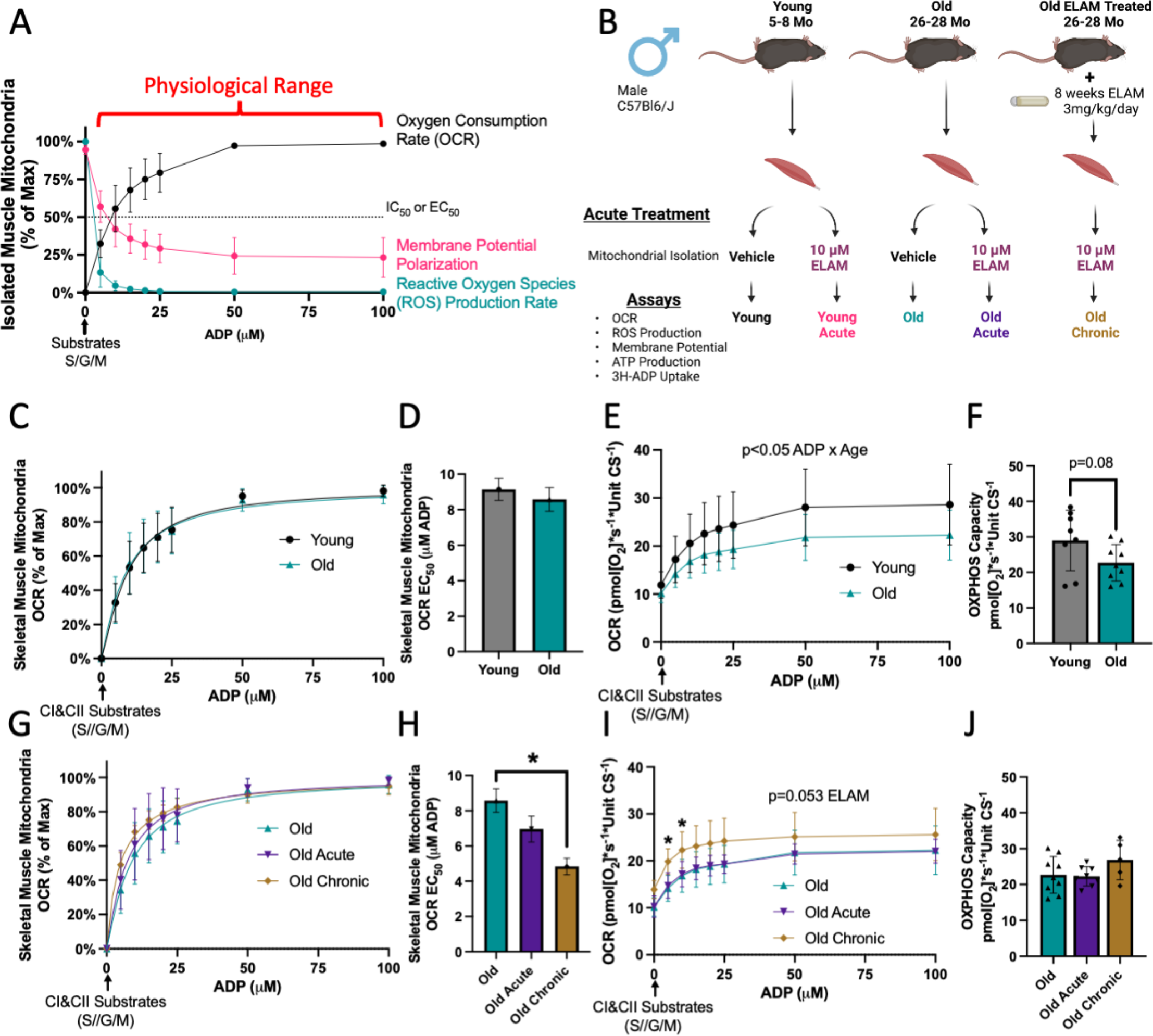
ELAM Improves Sensitivity to ADP for Respiration in Aged Muscle Mitochondria. A) Normalized representative traces of OCR, membrane potential polarization, and ROS production rate responses to ADP concentrations in 5 mo young, isolated muscle mitochondria. EC_50_ and IC_50_ are the concentrations of ADP required to stimulate or inhibit 50% of the response to ADP, respectively. B) Experimental design. C) Normalized OCR response to ADP, D) Calculated EC_50_, E) Raw OCR response to ADP, and F) Oxidative phosphorylation (OXPHOS) capacity from isolated muscle mitochondria of 5-8 mo young (n=8) and 26-28 mo old (n=9) male mice. G) Normalized OCR response to ADP, H) Calculated EC_50_, I) Raw OCR response to ADP, and J) OXPHOS capacity from isolated muscle mitochondria of 26-28 mo old (n=8-9) male mice with or without acute ELAM treatment and 26-28 mo old (n=5) male mice with chronic ELAM treatment. *p<0.05, **p<0.01 for post hoc tests or direct comparisons, main effect significant results are described in text and on graph. S/G/M (succinate, glutamate, malate respectively). Mean ± SD.

Using hexokinase clamp and exogenous cytochrome c, we observed no difference in OCR kinetic response to ADP titration with age in isolated mouse muscle mitochondria (**Figure 2C-D**). Across a range of ADP concentrations, total oxidative phosphorylation (OXPHOS) capacity was decreased in old mitochondria (p<0.05 ADP titration x Age effect by Two-Way RM ANOVA), and maximum respiration capacity trended towards a decrease (p=0.08 by t-test) (**Figure 2E-F**). Similar results were observed in muscle mitochondria from old female mice, i.e., OCR kinetic response to ADP was not different, total respiration was decreased across a range of ADP concentrations (p<0.05 by Two-Way RM ANOVA), and OXPHOS capacity was decreased significantly (p<0.05 by unpaired t-test) (**Figure S2A-D**). Overall, OXPHOS capacity was significantly decreased by age when including sex and age as variables (p<0.001 Age effect by Two-Way ANOVA).

Treatment with ELAM significantly increased sensitivity to ADP for respiration response in old chronic (8-week in vivo treatment) ELAM-treated mitochondria (p<0.01 by One-Way ANOVA and Tukey’s post hoc test) with a non-significant increase in ADP sensitivity in old acute-treated mitochondria by post hoc test (**Figure 2G-H**). Chronic ELAM treatment significantly increased respiration at low (≤ 10 μM) ADP concentrations (p<0.05 by Tukey’s post hoc test) and trended towards increased total respiration capacity (**Figure 2I-J**). Acute ELAM treatment had no effect in young isolated muscle mitochondria (**Figure S3A-M**).

To ensure this finding was not an artifact of reaction conditions, we performed the experiment with respiration buffer using only cytochrome c or only hexokinase clamp. Aging did not significantly affect ADP sensitivity in either condition, although ADP EC_50_ was increased ~12% in aged mitochondria with only cytochrome c (**Figure S4A-B, S4E-F**). Chronic ELAM treatment improved sensitivity to ADP in all buffer conditions (**Figure S4C-D, S4G-H**). In addition, acute ELAM significantly improved sensitivity to ADP in the absence of hexokinase clamp (**Figure S4D**).

We also tested ADP sensitivity for respiration in aged permeabilized muscle fibers, which is a more established model for comparing this phenotype (Holloway et al., 2018; Pharaoh et al., 2021). Isolated mitochondria were an order of magnitude more sensitive to ADP than permeabilized muscle fibers (**Figure S5A-B**). In permeabilized gastrocnemius fibers, acute and chronic ELAM treatment significantly increased sensitivity to ADP (**Figure S5A-B**). Total respiration was significantly decreased (p<0.0001 by two-way ANOVA) across a range of ADP concentrations with ELAM treatment; however, this was due to decreasing leak respiration without ADP (p<0.01 by Tukey’s posthoc test) (**Figure S5C**). OXPHOS coupling was not changed with treatment (**Figure S5D**). We have previously reported that acute (~1 hour) treatment also improved sensitivity to ADP in aged mouse muscle *in vivo* (Chavez et al., 2020). Acute treatment with ELAM significantly improved the kinetic response to ADP *in vivo*, in permeabilized muscle fibers, and in some isolated mitochondria conditions, which also demonstrates this phenotype is rapidly modifiable. Together, these data establish that ELAM is the first known pharmacological intervention that improves kinetic sensitivity to ADP in aged muscles from the level of isolated mitochondria up to *in vivo* conditions.

### ELAM Improves Membrane Potential Response to ADP in Aged Muscle Mitochondria

We used uptake of TMRM by isolated mitochondria to estimate relative mitochondrial membrane potentials. There was no difference in membrane potential response to ADP stimulation with age (**Figure 3A-C**). IC_50_ for membrane potential response was estimated to be less than 5 μM ADP for all groups, therefore our titration conditions did not have the sensitivity to identify the IC_50_ without large errors for membrane potential changes in response to ADP titration. Chronic treatment with ELAM significantly improved response to ADP in aged mitochondria for changes in membrane potential, both normalized as a proportion of the maximum polarization and as the total TMRM taken up (p<0.0001 ADP by treatment effect by Two-Way ANOVA) (**Figure 3D-E**). Chronic treatment with ELAM did not significantly affect maximum membrane potential polarization with substrates (**Figure 3F**). Overall, chronic ELAM treatment results in a more rapid response of membrane potential depolarization to ADP stimulation.

**Figure 3.**
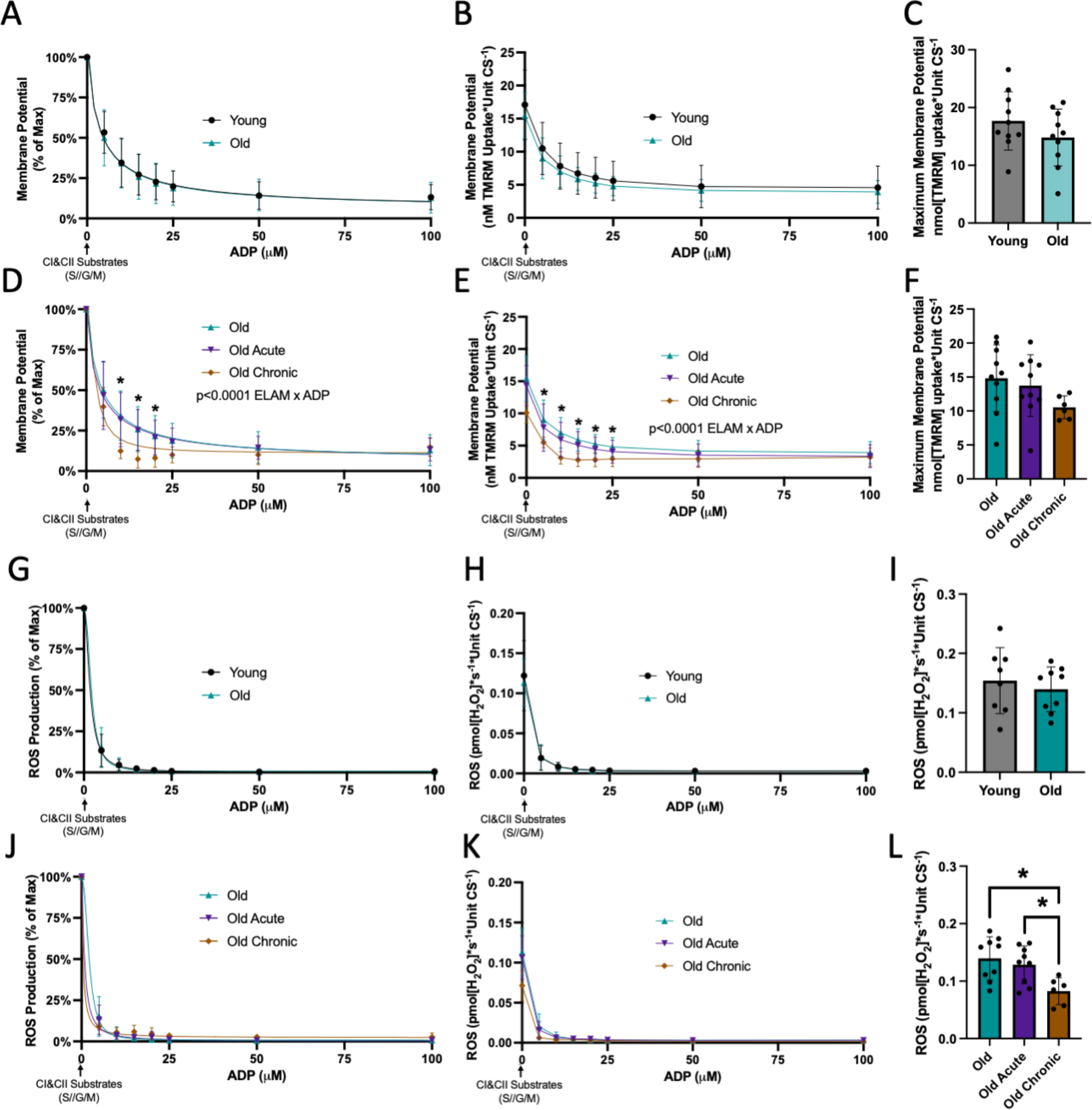
ELAM Increases Membrane Potential ADP Response and Decreases ROS Production. A) Normalized membrane potential response to ADP, B) Raw membrane potential response to ADP, and C) Maximum membrane potential from isolated muscle mitochondria of 5-8 mo young (n=10) and 26-28 mo old (n=10) male mice. D) Normalized membrane potential response to ADP, E) Raw membrane potential response to ADP, and F) Maximum membrane potential from isolated muscle mitochondria of 26-28 mo old (n=10) male mice with or without acute ELAM treatment and 26-28 mo old (n=6) male mice with chronic ELAM treatment. G)Normalized ROS production response to ADP, H) Raw ROS production response to ADP, and I) Maximum ROS production from isolated muscle mitochondria of 5-8 mo young (n=8) and 26-28 mo old (n=9) male mice. J) Normalized ROS production response to ADP, K) Raw ROS production response to ADP, and L) Maximum ROS production from isolated muscle mitochondria of 26-28 mo old (n=9) male mice with or without acute ELAM treatment and 26-28 mo old (n=6) male mice with chronic ELAM treatment. *p<0.05, **p<0.01 for post hoc tests or direct comparisons, main effect significant results are described in text and on graph. S/G/M (succinate, glutamate, malate respectively). Mean ± SD.

### ELAM Decreases Maximal ROS Production but Does Not Affect Kinetics in Aged Muscle Mitochondria

We measured ROS production in response to ADP stimulation in isolated mitochondria. There was no difference in ROS production with age in male or female muscle mitochondria (**Figure 3G-I, Figure S2E-G**). IC_50_ for ROS production was estimated to be less than 5 μM ADP for all groups, therefore our titration conditions did not have the sensitivity to identify the IC_50_ without large errors. Treatment with ELAM did not significantly affect the kinetic response of ROS production to ADP in aged mitochondria (**Figure 3J-K**). ROS production is highly stimulated after addition of the substrate succinate in the absence of ADP, a condition which results in the highest ROS production measured in our samples that tests the maximum limits of the samples to produce ROS. Chronic treatment with ELAM significantly decreased maximal ROS production in aged mitochondria (**Figure 3L**). In aged muscle fibers, chronic ELAM treatment did not affect kinetics, but significantly decreased ROS production across a range of ADP concentration (p<0.05 ELAM effect by Two-way RM ANOVA) (**Figure S5E-F**). Like isolated mitochondria, gastrocnemius fibers from chronic-treated ELAM mice trended towards decreased maximal ROS production (p = 0.055 by unpaired Welch’s t-test) (**Figure S5G**). Together, these data suggest chronic treatment with ELAM is required to impact ROS production in aged mitochondria oxidizing both complex I & II substrates.

### ELAM Increases Uptake of ADP Through the Adenine Nucleotide Translocator (ANT) and ATP Production in Aged Isolated Muscle Mitochondria

We measured uptake of [^3^H]ADP in young and old mitochondria from skeletal muscle isolated with vehicle or acute ELAM treatment. There was a significant effect of age on uptake of [^3^H]ADP uptake into the mitochondria; however, there was minimal difference between young and aged mitochondria in uptake in the physiological range of concentrations above 3 μM ADP (**Figure 4A**). Acute ELAM treatment increased uptake of [^3^H]ADP by mitochondria from aged skeletal muscle in nearly all samples and all ADP concentrations measured (p<0.05 ELAM effect by Two-Way RM ANOVA) (**Figure 4B-C**), but had no effect in young mitochondria (**Figure S3K**). We measured ANT specific ADP uptake by measuring uptake with or without the ANT inhibitor carboxyatractyloside (CAT) (**Figure S6**). This ANT-specific ADP transport accounted for approximately 80% of the total [^3^H]ADP uptake in this assay and was significantly increased with acute ELAM in old but not young muscle mitochondria (**Figure 4C, S3L, S6**).

**Figure 4.**
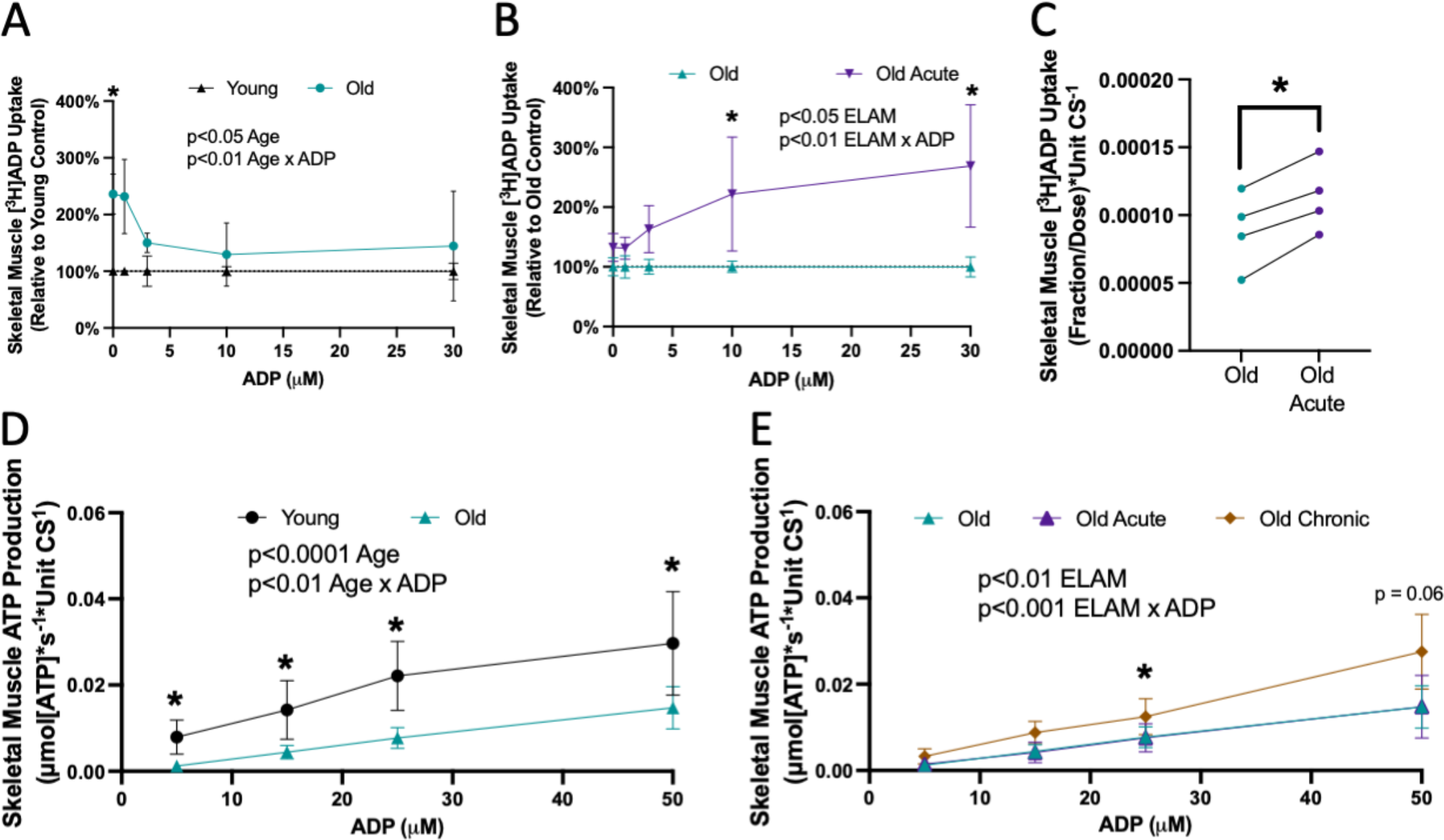
ELAM Increases Uptake of ADP Through the Adenine Nucleotide Translocator (ANT) in Aged Isolated Muscle Mitochondria and Restores ATP Production. A) Isolated mitochondria (n=3–5 per condition) from young and old skeletal muscle were incubated with a dose of [^3^H]ADP and increasing concentrations of ADP. The fraction of [^3^H]ADP dose was measured and used to estimate total ADP uptake under each ADP concentration and normalized to young control. B) Isolated mitochondria (n=3–5 per condition) from old and old acute ELAM-treated skeletal muscle were incubated with a dose of [^3^H]ADP and increasing concentrations of ADP. The fraction of [^3^H]ADP dose was measured and used to estimate total ADP uptake under each ADP concentration and normalized to old control. C) Isolated mitochondria (n=4 per condition) from old and old acute ELAM-treated skeletal muscle were incubated with a dose of [^3^H]ADP with or without 5 μM carboxyatractyloside (CAT) to inhibit ANT and calculate ANT-specific uptake of [^3^H]ADP. D) ATP production was measured in isolated mitochondria from 5-8 mo young (n=10) and 26-28 mo old (n=9) skeletal muscle across a range of ADP concentrations and normalized to ATP production without ADP. E) ATP production was measured in isolated mitochondria from 26-28 mo old (n=9) with or without acute ELAM treatment 26-28 mo old (n=5) or chronic ELAM treatment skeletal muscle across a range of ADP concentrations and normalized to ATP production without ADP. *p<0.05, **p<0.01 for post hoc tests, main effect significant results are described in text and on graph. Mean ± SD.

We measured ATP production in isolated mitochondria from skeletal muscle across a range of subsaturating ADP concentrations at equilibrium. ADP-stimulated ATP production was significantly decreased across all measured ADP concentrations in old muscle (**Figure 4D**). Chronic ELAM treatment significantly increased ATP production in aged muscle mitochondria (p<0.05 ELAM effect by matched mixed-effects model (REML) analysis) (**Figure 4E**). Acute treatment with ELAM did not affect ATP production in young or old muscle (**Figure 4E, S3M**). These data indicate that the improved response to ADP stimulation for membrane potential (**Figure 3D-E**) in aged mitochondria treated with ELAM for 8 weeks is being used to generate increased ATP production.

#### ELAM Increases Uptake of ADP Through the Adenine Nucleotide Translocator (ANT) in Aged Heart Isolated Mitochondria

To test if the effect of ELAM treatment on [^3^H]ADP uptake is specific to old muscle mitochondria, we measured uptake of [^3^H]ADP in young and old mitochondria from heart isolated with vehicle or acute ELAM treatment. There was a significant Age x ADP effect on the uptake of [^3^H]ADP into the heart mitochondria with an average of 35%-40% reduction in ADP uptake at 10-30 μM ADP (**Figure 5A**). Acute ELAM treatment significantly increased uptake of [^3^H]ADP by aged heart isolated mitochondria, specifically at the concentrations with the highest decrease in aged control samples (10-30 μM ADP) (**Figure 5B**). As in the muscle, ANT-specific ADP uptake accounted for approximately 80% of the total ADP transport and was significantly increased by ELAM treatment in old, but not young, heart mitochondria (**Figure 5C**).

**Figure 5.**
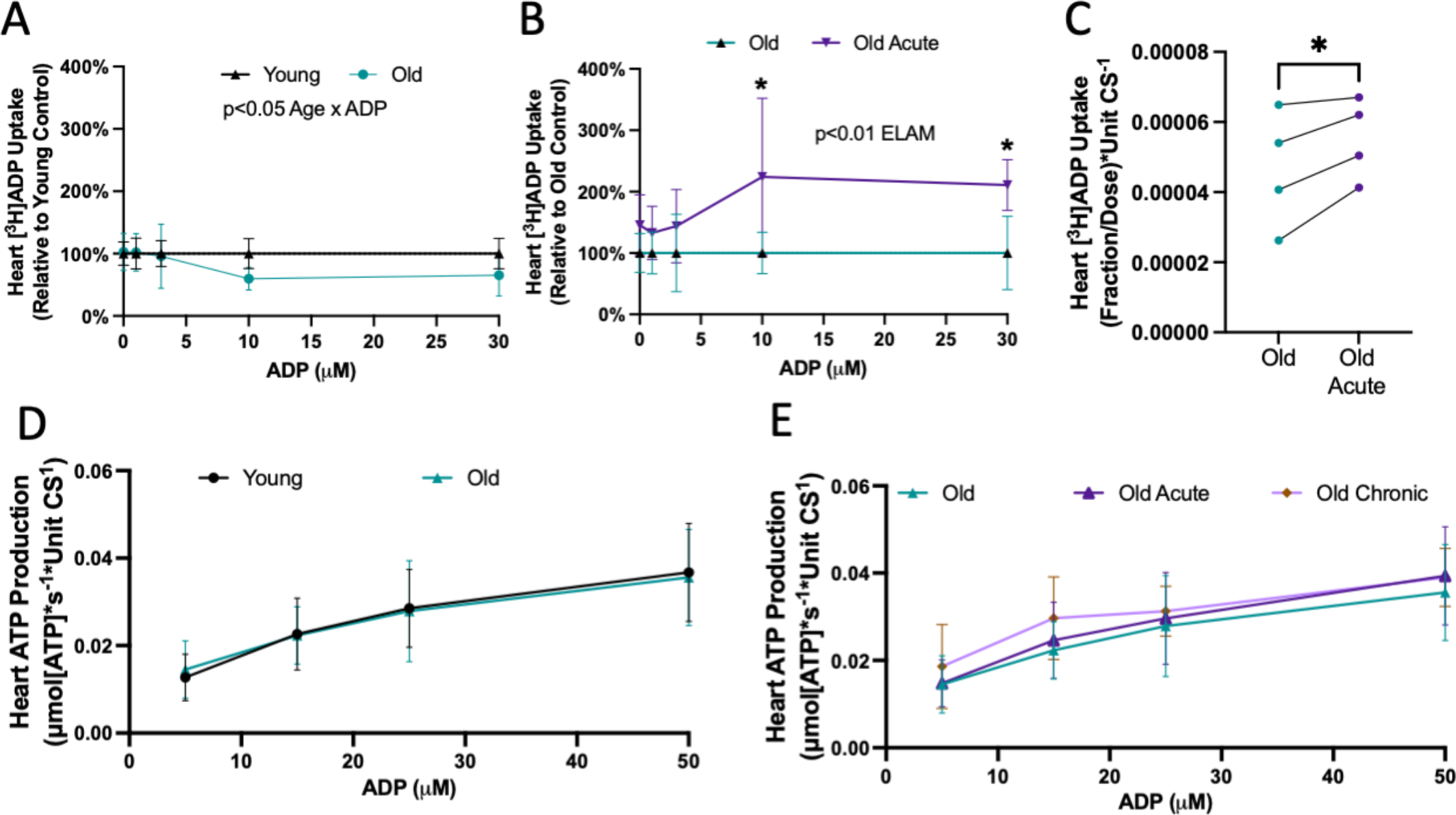
Acute ELAM Increases Uptake of ADP Through the Adenine Nucleotide Translocator (ANT) in Aged Heart Isolated Mitochondria. A) Isolated mitochondria (n=3–5 per condition) from young and old hearts were incubated with a dose of [^3^H]ADP and increasing concentrations of ADP. The fraction of [^3^H]ADP dose was measured and used to estimate total ADP uptake under each ADP concentration and normalized to young control B) Isolated mitochondria (n=3–5 per condition) from old and old acute ELAM-treated heart were incubated with a dose of [^3^H]ADP and increasing concentrations of ADP. The fraction of [^3^H]ADP dose was measured and used to estimate total ADP uptake under each ADP concentration and normalized to old control. C) Isolated mitochondria (n=4 per condition) from old and old acute ELAM-treated heart were incubated with a dose of [^3^H]ADP with or without 5 μM carboxyatractyloside (CAT) to inhibit ANT and calculate ANT-specific uptake of [^3^H]ADP. D) ATP production was measured in isolated mitochondria from 5-8 mo young (n=10) and 26-28 mo old (n=9) hearts across a range of ADP concentrations and normalized to ATP production without ADP. E) ATP production was measured in isolated mitochondria from 26-28 mo old (n=9) with or without acute ELAM treatment 26-28 mo old (n=5) or chronic ELAM treatment hearts across a range of ADP concentrations and normalized to ATP production without ADP. *p<0.05 for post hoc tests or paired t-test., main effect significant results are described in text and on graph. Mean ± SD.

We measured ATP production in isolated mitochondria from heart across a range of subsaturating ADP concentrations at equilibrium. ADP-stimulated ATP production was not affected by age or ELAM treatment in heart mitochondria (**Figure 5D-E**). These results suggest that acute effects of ELAM treatment on ADP uptake are separate from the effects on ATP production.

### Chronic ELAM Treatment Reverses Protein S-glutathionylation in the ADP/ATP Transport and Synthesis Pathways

Using mass spectrometry, we previously characterized changes in protein abundance, phosphorylation, and S-glutathionylation in muscle and heart from young, aged, and aged chronic ELAM treated mice (Campbell et al., 2019; Campbell et al., 2022; Chiao et al., 2020; Whitson et al., 2021). Here we reanalyzed and synthesized these datasets with a focus on proteins critical to ADP/ATP transport and ATP synthesis. There were no consistent changes with age or ELAM treatment on protein abundance or phosphorylation for isoforms of ANT, VDAC, creatine kinase, or ATP synthase in both skeletal muscle and heart tissues (**Figure S7A-B**). S-glutathionylation of cysteine residues significantly increased with age on several cysteines present on ANT, VDAC, creatine kinase, and ATP synthase proteins (**Figure 6A-C**). Treatment with ELAM significantly decreased S-glutathionylation of cysteine residues on several cysteines present on ANT, VDAC, creatine kinase, and ATP synthase proteins (**Figure 6A-C**). When comparing the change of all glutathionylated cysteines together, overall glutathionylation on each of these proteins increased with age whereas ELAM treatment had no or minimal difference from young in both aged skeletal muscle and heart (**Figure 6D-E**).

**Figure 6.**
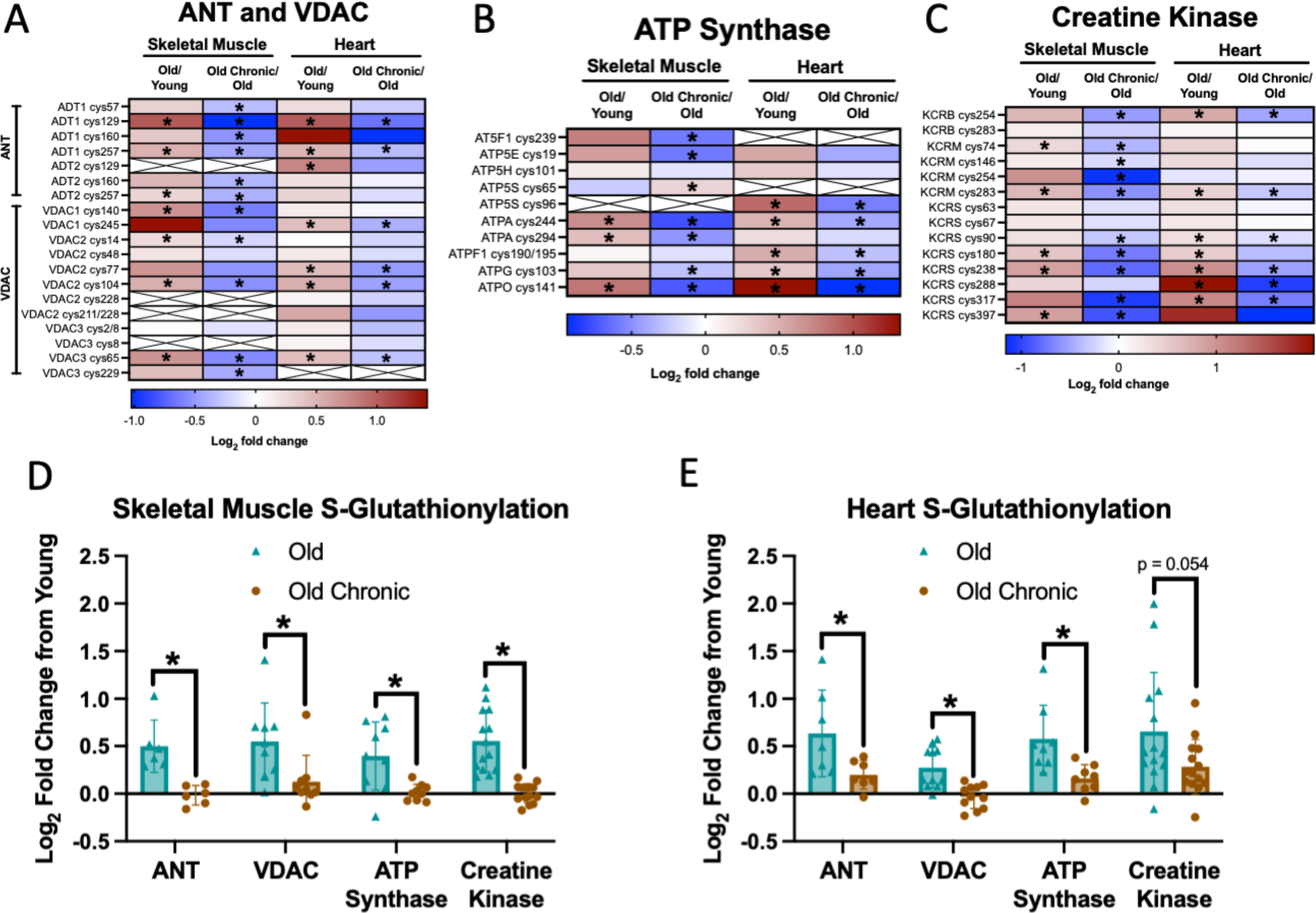
Chronic ELAM Treatment Reverses Protein S-Glutathionylation in the ADP/ATP Transport and Synthesis Pathway. Log_2_ fold change of s-glutathionylation of old/young and old with chronic ELAM treatment/old (n=5-6) in skeletal muscle and heart for A) ANT and VDAC proteins, B) ATP Synthase proteins, C) Creatine kinase proteins. Log_2_ fold change of s-glutathionylation of old/young and old with chronic ELAM treatment/young (n=5-6) at all sites for D) Skeletal muscle and E) Heart. *q<0.05 for each comparison in A-C. *p<0.05 for each comparison in D-E by t-test. Mean ± SD.

### ELAM Treatment Improves Aged Tissue Function in The Same Tissues it Improves Mitochondrial ADP Uptake

In the old mouse heart, systolic function decreased with age measured by global longitudinal strain (GLS), fractional shortening (FS), and ejection fraction (EF) (**Figure 7A-C, Supplemental Videos 1A-C**). Treatment with ELAM significantly improved systolic function, increasing GLS, FS, and EF in aged mice compared to pre-treatment values and restoring them to functional capacities comparable to young mice (**Figure 7E-G, Supplemental Videos 1A-C**). *In* vivo plantarflexor muscle force declined with age across a range of stimulation frequencies (**Figure 7D**). While treatment with ELAM did not increase maximum force production in this experiment, it significantly increased force production at lower frequency stimulations (**Figure 7H**). Together, ELAM treatment is associated with improved function in aged muscles that have increased mitochondrial [^3^H]ADP uptake, particularly in those that facilitate cardiac output.

**Figure 7.**
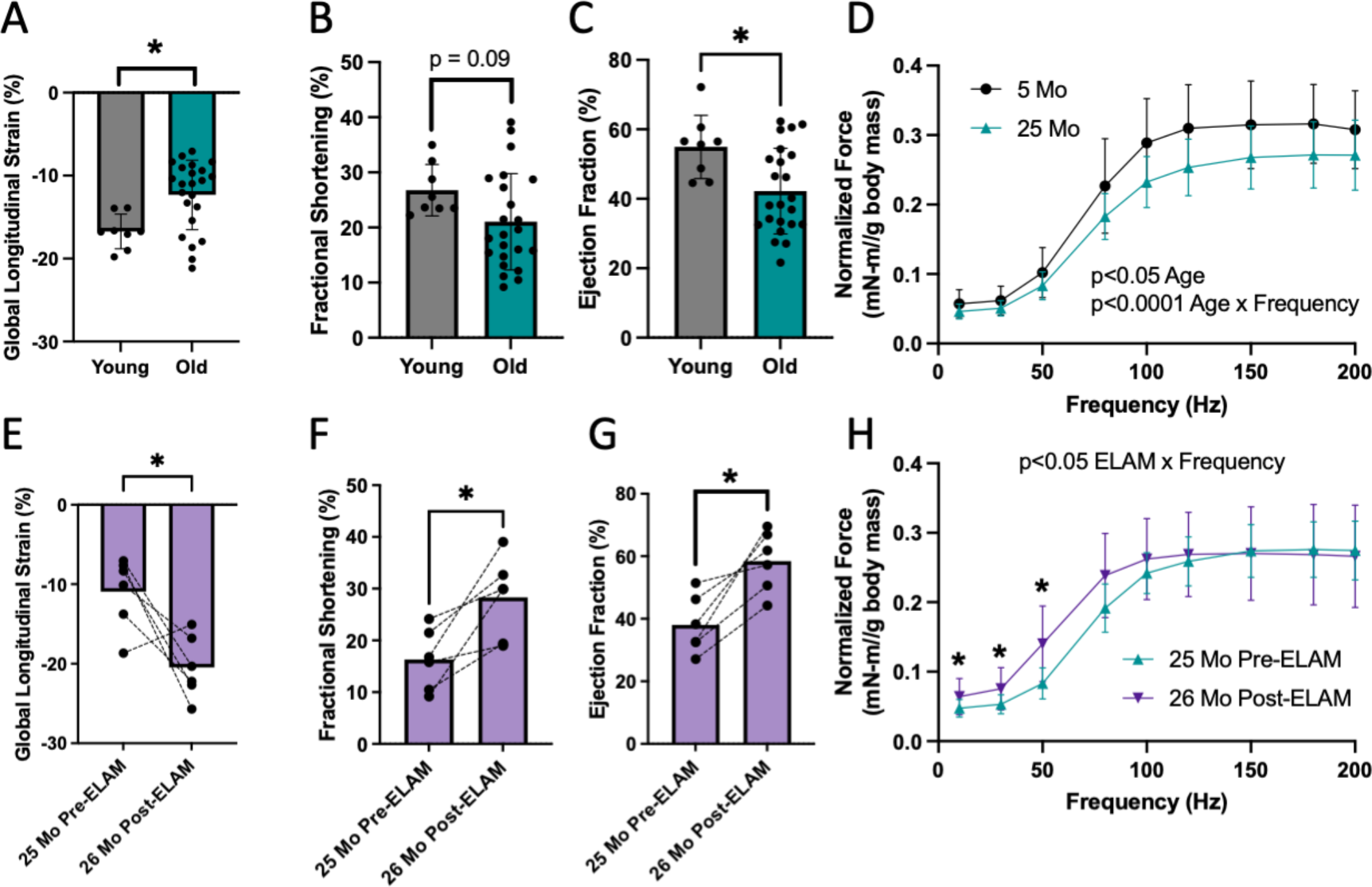
Treatment with ELAM Improves Physiological Function in Tissues with Improved ADP Uptake. Age comparison between young (5-8 mo) and old (25 mo) male mice for cardiac A) Global longitudinal strain (GLS), B) Fractional shortening (FS), C) Ejection fraction (EF), and D) *in vivo* hindlimb muscle force-frequency analysis normalized to body mass. Statistical comparisons in A-C by student’s t-test and in D by RM Two-Way ANOVA, *p<0.05, **p<0.01. Before and after chronic ELAM treatment comparison of old (25 mo) male mice for cardiac E) Global longitudinal strain (GLS), F) Fractional shortening (FS), G) Ejection fraction (EF), and for H) *in vivo* plantarflexor muscle force-frequency analysis normalized to body mass. Statistical comparisons in A-C by paired student’s t-test and in D by RM Two-Way ANOVA., *p<0.05, **p<0.01. Mean ± SD.

## DISCUSSION

### Major Findings

Here we expand on our previous *in vivo* report to demonstrate that ELAM improves sensitivity to ADP in old permeabilized muscle fibers and isolated muscle mitochondria (Chavez et al., 2020). Short term treatment with ELAM increased uptake of [^3^H]ADP by aged mitochondria in two muscle tissues, cardiac and skeletal muscle. We accounted for both ADP utilization at subsaturating conditions and the limitation of outer membrane damage by differential centrifugation to measure ADP response at subsaturaing conditions in isolated mitochondria. We use this technique to also measure ADP titration response for ATP production, OCR, ROS production, and membrane potential simultaneously in the same samples. Previous studies have been focused primarily on using respiration as a measure of ADP sensitivity. In aged skeletal and cardiac muscles, ELAM treatment improved ADP uptake and physiological function, providing further evidence for a link between the molecular and physiological phenotype.

### Current Progress on ADP Sensitivity in Muscle Research

Classic mitochondria experiments measure respiration at saturating conditions of ADP. These experiments determined capacities without measuring kinetics at more physiological conditions. Recently, kinetic experiments that measure response of mitochondria across a range of ADP concentrations have been reported that demonstrate loss of ADP sensitivity in aged muscle fibers from humans, mice, and rats (Gouspillou et al., 2014; Holloway et al., 2018; Pharaoh et al., 2021). Age-related ADP insensitivity has been associated with decreased OCR, decreased ATP production, and increased ROS production (Chavez et al., 2020; Gouspillou et al., 2014; Holloway et al., 2018; Pharaoh et al., 2021). In addition to aging, high fat diet, acute exercise, and oxidative stress have been reported to cause ADP insensitivity (Brunetta et al., 2021; Miotto et al., 2018; Pharaoh et al., 2021). However, despite being a major driver of aging muscle atrophy, loss of neuromuscular innervation does not affect muscle ADP sensitivity (Pharaoh et al., 2021). Mitochondrial ADP sensitivity is rapidly modifiable, although the mechanism underlying ADP insensitivity is currently unclear. Importantly, changes in protein content for VDAC, ANT, or other ADP/ATP exchangers were not sufficient to explain differences in ADP sensitivity across these studies.

It has been proposed that post-translational modifications of ANT likely modulate the rate of ATP/ADP transport (Barbeau et al., 2018; Miotto et al., 2018). ADP sensitivity is strongly affected by insulin signaling, and the decline in insulin sensitivity with age probably contributes to development of the phenotype (Brunetta et al., 2021; Miotto et al., 2018). ADP sensitivity is rapidly improved following insulin treatment in animals fed a high fat diet (Miotto et al., 2018). Exercise training has had mixed results with some studies finding increases in ADP sensitivity while others find decreases (Barbeau et al., 2018; Brunetta et al., 2021; Holloway et al., 2018; Miotto and Holloway, 2019; Ydfors et al., 2016). The exercise timing may explain the mixed results of exercise. Acute exercise decreases ADP sensitivity, but this is prevented by expression of the mitochondrial-targeted catalase (mCAT) antioxidant (Barbeau et al., 2018). In addition, ADP sensitivity was decreased in the *Sod1*^−/−^ mouse model of constitutive oxidative stress with accelerated sarcopenia (Pharaoh et al., 2021). Elevated mitochondrial ROS are a regulator of ADP sensitivity, likely by modification of ANT (Barbeau et al., 2018; Pharaoh et al., 2021).

### Aging ADP Sensitivity in Isolated Mouse Mitochondria

To our knowledge, this is the first research report to use isolated mouse muscle mitochondria as a model to assess ADP sensitivity in aging. Surprisingly, we did not observe a difference between young and old with hexokinase clamp and cytochrome c. It is possible that hexokinase clamp masked the effect of age on ADP sensitivity, since it decreased EC_50_ ~66%. In young and old isolated mitochondria with only cytochrome c, EC_50_ was non-significantly increased in old (12% increase) (**Figure S4B**). Although these results were not significant, the trend aligns with reports from human and mouse muscle fibers, mouse muscles *in vivo*, and rat isolated muscle mitochondria (Chavez et al., 2020; Gouspillou et al., 2014; Holloway et al., 2018; Pharaoh et al., 2021). Notably, in this reaction condition acute ELAM also significantly increases ADP sensitivity (**Figure S4D**). These results align with our findings in permeabilized muscle fibers, which demonstrate acute ELAM treatment improves ADP sensitivity in aged muscle fibers without cytochrome c or hexokinase clamp. (**Figure S5A-B**). ADP insensitivity is observed in old mice both in muscle fibers and *in vivo* but not in isolated mitochondria. This may be due to a limitation of isolated mitochondria from mice as a model for ADP sensitivity.

### Elamipretide is the First Known Pharmaceutical that Improves ADP sensitivity by Increasing ANT Transport

The rapid effects that acute exercise and insulin treatment exhibit on ADP sensitivity strongly suggest it is regulated at a post-translational level (Barbeau et al., 2018; Miotto et al., 2018). We have previously shown that one hour treatment with elamipretide improves muscle function, increases sensitivity to ADP, and decreases ADP concentration *in vivo* (Chavez et al., 2020; Siegel et al., 2013). We now extend this finding to show that ELAM rapidly improves muscle sensitivity to ADP *in vivo*, in permeabilized fibers, and in isolated mitochondria from aged mice. We previously reported the ADP concentration is ~30 μM ADP *in vivo* in resting aged mouse muscles (Siegel et al., 2013). The improved ADP uptake at these physiological ADP concentrations with acute ELAM treatment supports an important role for ANT function in the effect of ELAM on ADP sensitivity. One hour treatment with ELAM also decreased *in vivo* ADP concentration in resting aged muscle, consistent with increased uptake by the ANT (Siegel et al., 2013) and more efficient coupling of energy supply and demand under low energetic demand conditions. We propose two non-mutally exclusive mechanisms for this improved ANT function: direct interaction with the ANT protein and modification of ANT PTMs. We have previously reported that ELAM binds directly to ANT in the channel region of the protein (Chavez et al., 2020). Although the functional impact of this direct interaction has yet to be tested, the decrease in ANT mediated proton leak with acute ELAM treatment in aged cardiomyocytes suggests an impact on transport activity and selectivity (Zhang et al., 2020). Novel binding partners of ANT are an area of ongoing research interest due to its proposed role in mitochondrial dysfunction. In addition to classical ANT inhibitors, such as carboxyatractyloside and bongkrekic acid, it was recently identified that mitochondrial uncouplers interact directly with ANT to induce proton leak {Bertholet, 2022 #294}. In addition to the direct binding hypothesis, here we report that ELAM treatment decreases s-glutathionylation of ANT that occurs with age and increases ADP uptake through the ANT. ELAM increasing sensitivity to ADP through an effect on redox sensitive PTMs as a mechanism of action would explain why ELAM effectively increases ADP sensitivity in aging and the *Sod1*_−/−_ mouse but does not affect young mitochondria (Pharaoh et al., 2021).

Surprisingly, acute ELAM increased uptake of ADP through the ANT in isolated mitochondria but did not increase ATP synthesis; however, chronic ELAM treatment increased ATP production in aged muscle. The increase in resting ADP concentration and decrease in ATP concentration with age is consistent with deficiencies in ADP import or ATP production (Campbell et al., 2019; Siegel et al., 2013). Both short term (1 hour) and long term (8 week) treatment with ELAM decreased ADP concentration in muscle and increased maximal ATP production, suggesting that the effects of ELAM treatment in isolated mitochondria reported here translate to *in vivo* muscle function as well (Campbell et al., 2019; Siegel et al., 2013).

### Elamipretide Improves Tissue Function in Aged Tissues with Increased Mitochondrial ADP Uptake

In this paper we demonstrated that ELAM treatment repaired muscle force at submaximal tetanus and restored heart systolic function, both of which exhibited increased mitochondrial uptake of ADP with ELAM treatment. Our aging results agree with recent publications in mice that identified a longitudinal decline with age in cardiac GLS and EF using strain echocardiography, which established the aging mouse as a model for decreased GLS (de Lucia et al., 2019). GLS and EF using strain echocardiography have recently been reported as improved methods to measure systolic dysfunction in humans (Karlsen et al., 2019; Verdonschot et al., 2021). Here for the first time, we demonstrate reversal of age-related declines of GLS in mice and restoration of systolic function using ELAM treatment. This improved systolic function adds to our previous reports that ELAM treatment improves diastolic function in old mice (Chiao et al., 2020; Whitson et al., 2020). In addition, increased *in vivo* muscle force at submaximal stimuli in aged mice after ELAM treatment builds on previous reports of increased muscle *in vivo* maximal ATP production, fatigue resistance, and exercise capacity (Campbell et al., 2019; Siegel et al., 2013). Some improvements in both cardiac function and muscle strength were also reported with ELAM treatment in patients with Barth Syndrome, an ultrarare genetic condition caused by abnormal and dysfunctional cardiolipin on the inner mitochondrial membrane (Reid Thompson et al., 2021). Together, these data support the hypothesis that ELAM treatment improves ADP sensitivity and muscle function, and provide further evidence that age-related ADP insensitivity plays a role in aging muscle weakness.

### Limitations

Several limitations occurred in this study. The Oroboros O2k is a highly sensitive instrument, however it has low throughput. Due to limitations in the number of chambers, we could only measure simultaneous OCR, ROS, and membrane potential in mitochondria from one tissue and did not make measurements in the hearts from these animals. With hexokinase clamp and cytochrome c, the EC_50_ and IC_50_ for mitochondrial responses to ADP were less than 10 μM ADP, an order of magnitude lower than for values for muscle fibers or rat isolated mitochondria with only hexokinase clamp (Gouspillou et al., 2014). The lowest titration of ADP used in this set of experiments was 5 μM ADP, so future experiments with isolated mitochondria will include more low-dose ADP titrations to increase precision. In addition, due to the experimental complexity, cost, and amount of mitochondria required, we were unable to measure [^3^H]ADP uptake in chronically treated heart or muscle mitochondria. Finally, the majority of the data reported in this manuscript are from male mice; however, we have previously reported beneficial effects of ELAM treatment on mitochondrial function and heart and muscle physiological function in female mice (Campbell et al., 2019; Chiao et al., 2020; Siegel et al., 2013).

### Conclusions and Future Directions

ELAM treatment increases sensitivity to ADP for old muscle *in vivo*, in permeabilized fibers, and in isolated mitochondria, acutely increases uptake of ADP into aged mitochondria through the ANT, decreases S-glutathionylation of the ANT, and repairs whole tissue function. Future directions will focus on identifying the effects of direct ELAM binding to ANT and how modulating specific PTMs in ANT affect ADP sensitivity and tissue function.

## METHODS

### Animals

C57Bl6/J male and female mice were acquired from The National Institute on Aging (NIA) mouse colony and housed at the University of Washington in a specific-pathogen free facility. All mice were maintained at 21 °C on a 14/10 light/dark cycle at at 30-70% humidity and given standard mouse chow (LabDiet PicoLab^®^ Rodent Diet 20) and water *ad libitum* with no deviation prior to or after experimental procedures. This study was reviewed and approved by the University of Washington Institutional Animal Care and Use Committee (IACUC).

### Chronic 8-Week Elamipretide Treatment

Aged mice were treated with ELAM by osmotic minipumps for 8 weeks as previously described (Campbell et al., 2019). *In vivo* muscle force and echocardiography were measured in the week prior to pump implantation. The pumps were implanted in 25-month old male mice and were replaced after 4 weeks. Post treatment muscle force and echocardiography were measured 6-7 weeks after initial pump implantation. Mice were euthanized and tissues collected for mitochondrial isolation at 7-8 weeks treatment at ~27 months old. ELAM was provided by Stealth BioTherapeutics Inc.

### Mitochondrial Isolation and Acute ELAM Treatment

The gastrocnemius, quadriceps femoris, and tibialis anterior (TA) muscles from both hindlimbs and the heart were dissected from young (5–8 mo) and old (26–28 mo) mice, and mitochondrial isolation was performed by differential centrifugation. The tissues were homogenized using a high-speed drill on ice in a glass Dounce homogenizer in Mitochondria Isolation Buffer (containing in mM 210 sucrose, 2 EGTA, 40 NaCl, 30 HEPES, pH 7.4). The homogenate was centrifuged at 900 × g at 4 °C for 10 minutes. The supernatant was collected and centrifuged at 10,000 × g at 4 °C for 10 minutes. The supernatant was removed, and the mitochondrial pellet was resuspended in ice-cold Respiration Buffer (RB) without bovine serum albumin (BSA) (1.5 mM EGTA, 3 mM MgCl_2_-6H_2_O, 10 mM KH_2_PO_4_, 20 mM HEPES, 110 mM sucrose, 100 mM mannitol, 60 mM K-MES, 20 mM taurine, pH 7.1). For acute and chronic ELAM treatments, one set of hindlimb muscles and half of the heart were isolated with vehicle Mitochondria Isolation Buffer, and the other set of muscles and other half of the heart were isolated with Mitochondria Isolation Buffer with 10 μM ELAM (**Figure 2B**). The mitochondrial pellets were resuspended in vehicle RB without BSA or RB without BSA +10 μM ELAM. Isolated mitochondria protein concentration was determined using standard Bradford Assay procedures.

### Simultaneous Oxygen Consumption Rate, Reactive Oxygen Species Production, and Membrane Potential

Mitochondrial respiration, ROS production, and membrane potential in response to an ADP titration were assayed in isolated skeletal muscle mitochondria using an Oxygraph 2K dual respirometer/fluorometer (Oroboros Instruments, Innsbruck, Austria) (**Figure 2A**). RB with taurine and BSA was used for respiration measurements (1.5 mM EGTA, 3 mM MgCl_2_-6H_2_O, 10 mM KH_2_PO_4_, 20 mM HEPES, 110 mM Sucrose, 100 mM Mannitol, 60 mM K-MES, 20 mM taurine, 1 g/L BSA, pH 7.1). Hexokinase clamp (1 U/mL hexokinase, 2.5 mM D-glucose) was used to maintain equilibrium of ATP/ADP at subsaturating ADP concentrations (**Figure 1A**) (Lark et al., 2016). To measure ROS production, 10 μM Amplex UltraRed, 1 U/mL horseradish peroxidase (HRP), and 5 U/mL superoxide dismutase (SOD) were added to the chamber. A hydrogen peroxide standard curve was performed each day and used to convert fluorescence signal to ROS concentration. Membrane potential was measured by adding 1 μM tetramethylrhodamine, methyl ester (TMRM) to the chamber. Respirometry and fluorometry reagent stocks were prepared according to Oroboros’s instructions (bioblast.at). Respiration was measured at 37°C with stirring during substrate and inhibitor titrations, and the chambers were hyper-oxygenated to 450-500 μM O_2_.

For the substrate-uncoupler-inhibitor titration (SUIT) protocol, first 10 μM ELAM was added to the acute treatment chambers. Next, 10 μM cytochrome c was added to each chamber to allow measurement of respiration in isolated mitochondria without limitation by membrane damage occurring during isolation (**Figure 1A**). Approximately 100 μg mitochondrial homogenate was added to each 2 mL chamber. CI&CII-linked leak respiration was stimulated by adding 10 mM succinate, 10 mM glutamate, and 0.5 mM malate. ADP stimulated respiration was measured during a titration of ADP from 0-2000 μM ADP. For final OCR values after each titration, the OCR was calculated by taking the average value from the chamber with Amplex UltraRed and the chamber with TMRM for each sample. The background OCR with de-energized mitochondria was subtracted from all measured functional parameters before reporting final values.

A quality control cutoff was applied to all samples requiring OXPHOS coupling higher than 0.25 (average±SD = 0.49±0.12, cutoff is >2 SD from mean) or Vmax of respiration at ADP concentrations higher than 10 μM (average±SD = 95±5.7, cutoff >14 SD from mean). Both cutoffs indicate a lack of respiration response to ADP titration, which affects the curve fitting used to calculate the kinetics of ADP response. Using this cutoff, two young and two old samples were removed from analysis.

Previously, ADP sensitivity studies have used the Michaelis-Menten (MM) to fit the OCR response to ADP titration and calculate Km. We compared MM curve fit to [Agonist] vs normalized dose response with Graphpad Prism [Agonist] vs response had the better fit for all groups and was the preferred model due to having a higher r-squared for each data set. The settings for the [Agonist] vs. normalized response -- Variable slope equation were: Detect and eliminate outliers at q=1%, Fitting method - least squares, Convergence criteria – medium, no weighting, replicates - consider each replicate y values as an individual point.

### Citrate Synthase Activity Assay

Citrate Synthase (CS) activity is reportedly a more accurate marker of mitochondrial mass than total protein content when performing comparisons across age (Figueiredo et al., 2008). CS activity assay was performed on mitochondrial isolations and used to normalize mitochondrial respiration. CS Activity was measured by spectrometric quantitation (412 nm) of 5,5’dithiobis-2-nitrobenzoic acid conversion to 2-nitro-5-thiobenzoic acid in the presence of Coenzyme A thiol generated during citrate production (CS0720, Sigma) as previously described (Crouch et al., 2017).

### [^3^H]ADP Uptake

Uptake of [^3^H]ADP was measured in isolated muscle and heart mitochondria. Radioactive [^3^H]ADP (American Radiolabeled Chemicals, 1 mCi/mL) was diluted in RB+BSA to approximately 0.2 μCi per 10 μL. Measurements of mitochondrial uptake of [^3^H]ADP were carried out as previously described (Sweet et al., 2004). Microcentrifuge tubes were placed in a water bath heated to 37°C and reaction combinations were added to a total volume of 190 μL including buffer (RB+BSA with or without 10 μM ELAM) and mitochondria (~40-50 μg per reaction). Some conditions were performed with 1-100 μM [^1^H]ADP (Sigma A 5285) or with 5 μM carboxyatractyloside (Cayman Chemical Item No. 21120) to inhibit ANT (**Figure 1A, S6**). Each experimental condition received 10 μL of the [^3^H]ADP dose to bring the final reaction volume to 200 μL which initiated radioactive ADP uptake. After 1 min, the mitochondria-associated radioactivity was transferred to 0.4-mL centrifuge tubes (USA Scientific, Ocala, FL) containing a 100-uL layer of oil consisting of 1:37.5 n-dodecane:bromododecane (Sigma) and placed into a Beckman E centrifuge. At exactly 90 seconds after addition of [^3^H]ADP the tubes were centrifuged at 12,535⍰g for 5 seconds thereby separating the mitochondria-associated radioactivity from the free. The centrifuge tubes were flash frozen in liquid nitrogen and the bottom of the microcentrifuge tube containing the mitochondrial pellet was cut off and placed into a glass scintillation vial. Vials were subsequently filled with 5 mL scintillation fluid (Ecolume, MP Biomedicals). All samples were counted usage a Beckman Liquid Scintillation Counter (Model LS6500) and the fraction of the total [^3^H]ADP dose present in the mitochondrial fraction was calculated. All values were corrected for non-specific background by subtracting count obtained from samples containing the same [^3^H]ADP concentration, 100 μM standard ADP buffer, and mitochondria, but where the reaction was initiated on ice and immediately spun down. [^1^H]ADP competes with [^3^H]ADP for uptake by the ANT. ADP uptake was measured over a range of [^1^H]ADP concentrations with the same dose of [^3^H]ADP. As the total concentration of ADP increased, the fraction of the [^3^H]ADP dose taken up by mitochondria during the assay decreased. To ease data interpretability across the range of ADP concentrations, the values were normalized to the control group at each ADP concentration to compare ADP uptake rates between groups at each total ADP concentration.

### ATP Production

ATP Production rates were measured in muscle and heart isolated mitochondria using an extended version of the hexokinase clamp that also includes NADP+ and glucose-6-phosphate dehydrogenase (G-6-PDH) modified from a previous publication (**Fig. 1A**) (Lark et al., 2016). The ATP produced by the mitochondria is used to produce NADPH in a 1:1 stoichiometry, and the concentration of NADPH is estimated using an NADPH standard curve (MP Biomedicals 2646-71-1) (**Figure 1A**). The reaction was modified to a high throughput 96-well plate format allowing many measurements in parallel. Reaction buffer was prepared with identical conditions to the respiration experiments and G-6-PDH and NADP+. The reaction buffer consisted of RB+BSA with 1 U/ml Hexokinase, 2.5 mM D-glucose, 2.5 U/ml G-6-PDH, 2.5 mM NADP+, 10 mM Succinate, 10 mM glutamate, 0.5 mM malate, and 10 μM cytochrome c (all reagents from Sigma). Buffer was prepared with or without 10 μM ELAM and pre-warmed to 37°C in a water bath. NADPH standard curve or 0-50 μM ADP concentrations were added to wells in a black 96-well plate and pre-warmed to 37°C in an incubator. Isolated mitochondria were diluted to 0.05 mg protein/mL in RBA+BSA with or without 10 μM ELAM. 100 μL of reaction buffer was added to each well, and the reaction was initiated by adding 100 μL of mitochondria to each well, bringing the final concentration to 25 μg/ml (5 μg per well). The plate was immediately loaded into a Biotek Synergy 4 96-well plate reader (Agilent Technologies, USA), and NADPH fluorescence (340/460 excitation/emission) was measured every 2 minutes for 30 minutes with continuous shaking between reads. The fluorescent signal was converted to NADPH concentration using an NADPH standard curve measured simultaneously on the plate at each time point. The increase in NADPH concentration over 10 minutes was calculated, converted 1:1 to ATP, and normalized to 0 μM ADP to analyze ATP production in response to ADP stimulation.

### Muscle Fiber Preparation and Respirometry and Fluorometry

Permeabilized red gastrocnemius muscle fiber bundles were prepared as previously described with some modifications (Ahn et al., 2018; Kuznetsov et al., 2008; Pharaoh et al., 2021). Briefly, a small piece of red gastrocnemius muscle was dissected and ~3-5 mg fiber bundles were separated in ice-cold isolation buffer (IB containing (in mM) 7.23 K_2_EGTA, 2.77 CaK_2_EGTA, 20 imidazole, 0.5 DTT, 20 taurine, 5.8 ATP, 15 PCr, 6.6 MgCl_2_–6H_2_O, 50 K-MES, pH 7.1). The fiber bundles were permeabilized for 40 minutes in ice-cold IB with 50 μg/mL saponin for 40 minutes, then washed in ice-cold IB for 5 minutes and respiration buffer (RB containing (in mM) 110 sucrose, 100 mannitol, 60 K-MES, 30 KCl, 20 HEPES, 20 taurine, 10 K_2_HPO_4_, 3 MgCl_2_–6H_2_O, 1.5 EGTA, 1 mg/mL Bovine Serum Albumin (BSA), pH 7.1) once for 5 minutes and once for 15 minutes. Fibers were permeabilized and washed at each step with vehicle or vehicle + 10 μM ELAM.

Assay conditions and explanation of methods for simultaneous measurement of OCR using the Oroboros Oxygraph-2k (O2k, OROBOROS Instruments, Innsbruck, Austria) and hydroperoxide production rate using the O2k-Fluo LED2-Module Fluorescence-Sensor Green with Amplex UltraRed were previously described (Ahn et al., 2018; Kuznetsov et al., 2008; Pharaoh et al., 2021). For CI&CII-linked respiration and ROS Production, the samples were measured in RB at 37 °C with 10 μM Amplex UltraRed, 1 U/mL HRP, and 5 U/mL SOD. Vehicle or 10 μM ELAM were added to the chambers after calibration, then the chambers were hyper-oxygenated to 450-500 μM O_2_. Fiber bundles were added, then Leak respiration was induced with 10 mM succinate, 5 mM pyruvate, 10 mM glutamate, and 0.5 mM malate. ADP was titrated (0-6,000 μM) to induce respiration. Next, 0.5 μM rotenone was added to measure CII-linked respiration. Finally, 2.5 μM antimycin A was added to measure non-mitochondrial respiration, and all other values were normalized using this value to remove OCR not from the electron transport chain. The fluorescent Amplex UltraRed signal was converted to hydroperoxide via a standard curve established on each day of experiments. Background resorufin production was subtracted from each measurement. Data for both OCR and rates of hydroperoxide generation were normalized by milligrams of muscle bundle wet weights.

### Abundance, Phosphorylation, and Redox Proteomics

Methods and the full datasets for abundance, phosphorylation, and redox proteomics were previously reported (Campbell et al., 2019; Campbell et al., 2022; Chiao et al., 2020; Whitson et al., 2021). We have analyzed a subset of these datasets here focused on ADP/ATP pathways.

### Echocardiography Imaging and Analysis for Cardiovascular Function

2D echocardiography of the left ventricle (LV) was performed using the Vevo 3100 Preclinical Imaging System, Vevo Imaging Station, and MS400 probe from VisualSonics (Toronto, Canada). Mice are anesthetized with 4% isoflurane and fur is removed from their chest with Nair. The mice are placed in a supine position and heart rate, respiration rate, and core body temperature are monitored using the Vevo Animal Monitoring System SM200 and Vevo Monitor App software from VisualSonics. The SM200 also provides heat support to the mice while they are maintained on 1-3% isoflurane anesthesia. Parasternal long axis view (PLAX) B-mode and parasternal short-axis view (PSAX) B-mode and M-mode images were acquired. Resting cardiac workload images were acquired while maintaining the heart rate of the mouse between 400-500 beats per minute (bpm). Echocardiography was analyzed using the Vevo LAB software with Vevo Strain package. Strain analysis, including global longitudinal strain (GLS), was analyzed from 5 consecutive cardiac cycles without a respiration from the PLAX B-mode images using Vevo Strain package. The results were exported to Microsoft Excel and statistical analysis and graphing was performed using GraphPad Prism software. The age comparisons used values from young (8 mo) and old (25 mo) mice including the baselines values for the ELAM chronic treatment mice.

### In Vivo Muscle Force

*In vivo* measurement of muscle force was performed as previously described in the hindlimb plantarflexor muscles (Valencia et al., 2021). The age comparisons used values from young (8 mo) and old (25 mo) mice including the baselines values for the ELAM chronic treatment mice.

### Statistics and graphing

Graphing and statistical analysis were performed using Microsoft Office Excel and GraphPad Prism for OS X (GraphPad Software, San Diego, California USA). For all statistical tests, alpha levels were set to p<0.05. *p<0.05 for post hoc tests or direct comparisons unless described otherwise. IC_50_ for ROS and membrane potential in isolated mitochondria was determined using nonlinear regression [Inhibitor] vs. Response – variable slope (four parameters) equation, and EC_50_ for OCR was determined using [Agonist] vs normalized dose response. For comparisons across a range of ADP concentrations or frequencies for muscle contraction, young vs old, young vs young acute, and old vs old acute were compared by two-way RM ANOVA with Sidak’s post hoc test. Old vs old acute vs old chronic were compared with mixed-effects model (REML) with Tukey’s post hoc test. For heart function, pre- and post-ELAM treatment was compared by paired two-tailed student’s t-test. Gastrocnemius fibers ROS production was compared by unpaired Welch’s t-test, because the variances were significantly different. Principal component analysis was performed using ClustVis as previously described (Metsalu and Vilo, 2015; Pharaoh et al., 2019). Main effect significant results are described in text and on graph. Plots depict mean ± standard deviation.

## Acknowledgments

ELAM was provided by Stealth BioTherapeutics Inc. at no cost. The authors would like to thank Ana Valencia, Amy Martinson, and the University of Washington Center for Cardiovascular Biology for training and assistance on experimental methods.

## Author Contributions

GP and DJM created the study design. GP, VK, RSS, JW, MMP, and MDC conducted experiments. GP, VK, SK, RSS, MMP, MDC analyzed experimental data. WJQ, MJM, JV, PR, IRS, DJM oversaw experiments and provided laboratory resources. GP and DJM wrote and edited the manuscript. All authors edited and prepared the final manuscript for submission.

