## Supplemental Information for "Elamipretide Improves ADP Sensitivity in Aged Mitochondria by Increasing Uptake through the Adenine Nucleotide Translocator (ANT)"

**SUPPLEMENTAL FIGURES**

**Figure S1. Comparisons of Multivariate Mitochondrial Data**.

A) Principal component analysis (PCA) using all available mitochondrial data from each sample.

B) Comparison of membrane potential and OCR for each sample during leak state after addition of succinate and succinate, glutamate, and malate before titration of ADP.

C) Comparison of OCR and ROS for each sample during leak state after addition of succinate and succinate, glutamate, and malate before titration of ADP.

D) Comparison of membrane potential and ROS for each sample during leak state after addition of succinate and succinate, glutamate, and malate before titration of ADP.

**Figure S2. ADP Kinetic Sensitivity is Not Changed but OXPHOS Capacity is Reduced with Age in Isolated Female Muscle Mitochondria.**

A) Normalized OCR response to ADP,

B) Calculated EC_50_,

C) Raw OCR response to ADP, and

D) OXPHOS capacity from isolated gastrocnemius muscle mitochondria of 6 mo young (n=3) and 30 mo old (n=3) female mice.

E) Normalized ROS production response to ADP,

F) Raw ROS production response to ADP, and

G) Maximum ROS production from isolated gastrocnemius muscle mitochondria of 6 mo young (n=3) and 30 mo old (n=3) female mice.

*p<0.05 for post hoc tests or direct comparisons, main effect significant results are described in text and on graph. S/G/M (succinate, glutamate, malate respectively). Mean ± SD.

**Figure S3. Acute ELAM Treatment Does Not Affect Young Muscle Mitochondria.**

A) Normalized OCR response to ADP,

B) Calculated EC_50_,

C) Raw OCR response to ADP,

D) OXPHOS capacity,

E) Normalized membrane potential response to ADP,

F) Raw membrane potential response to ADP,

G) Maximum membrane potential,

H) Normalized ROS production response to ADP,

I) Raw ROS production response to ADP, and

J) Maximum ROS production from isolated muscle mitochondria of 5-8 mo young (n=8) male mice with or without acute ELAM treatment.

K) Isolated mitochondria (n=3–5 per condition) from young skeletal muscle with vehicle or acute ELAM treatment were incubated with a dose of [^3^H]ADP and increasing concentrations of ADP. The fraction of [^3^H]ADP dose was measured and used to estimate total ADP uptake under each ADP concentration and normalized to young control.

L) Isolated mitochondria (n=3–5 per condition) from young skeletal muscle with vehicle or acute ELAM treatment were incubated with a dose of [^3^H]ADP with or without 5 µM carboxyatractyloside (CAT) to inhibit ANT and calculate ANT-specific uptake of [^3^H]ADP.

M) ATP production was measured in isolated mitochondria from young skeletal muscle with vehicle or acute ELAM treatment across a range of ADP concentrations and normalized to ATP production without ADP.

Mean ± SD.

**Figure S4. ELAM Improves ADP Sensitivity for Oxygen Consumption Rate in Aged Isolated Mitochondria Across all Buffer Conditions.**

A) OCR response curve to ADP, and

B) Calculated EC_50_ in isolated muscle mitochondria with only 10 µM exogenous cytochrome c (no hexokinase clamp) from isolated muscle mitochondria of 5-8 mo young (n=4) and 26-28 mo old (n=4) male mice.

C) OCR response curve to ADP, and

D) Calculated EC_50_ in isolated muscle mitochondria with only 10 µM exogenous cytochrome c (no hexokinase clamp) from isolated muscle mitochondria of 26-28 mo old (n=4) male mice with or without acute ELAM treatment and 26-28 mo old (n=6) male mice with chronic ELAM treatment.

E) OCR response curve to ADP, and

F) Calculated EC_50_ in isolated muscle mitochondria with only hexokinase clamp (no exogenous cytochrome c) from isolated muscle mitochondria of 5-8 mo young (n=4) and 26-28 mo old (n=4) male mice.

G) OCR response curve to ADP, and

H) Calculated EC_50_ in isolated muscle mitochondria with only hexokinase clamp (no exogenous cytochrome c) from isolated muscle mitochondria of 26-28 mo old (n=4) male mice with or without acute ELAM treatment and 26-28 mo old (n=6) male mice with chronic ELAM treatment.

Significance determined by one-way ANOVA with Tukey’s posthoc test, *p<0.05. Mean ± SD.

**Figure S5. ELAM Effects on ADP Sensitivity are Replicated in Aged Muscle Fibers.**

A) Normalized OCR response to ADP,

B) Calculated EC_50_,

C) Raw OCR response to ADP,

D) OXPHOS coupling,

E) Normalized ROS production response to ADP,

F) Raw ROS production response to ADP, and

G) Maximum ROS production from isolated muscle mitochondria of 26-28 mo old (n=5) male mice without acute ELAM treatment and 26-28 mo old (n=4) male mice with chronic ELAM treatment.

*p<0.05 for post hoc tests or direct comparisons, main effect significant results are described in text and on graph. S/P/G/M (succinate, pyruvate, glutamate, malate respectively). Mean ± SD.

**Figure S6. Proportion of Total [^3^H]ADP Uptake with and without Carboxyatractyloside (CAT).**

Isolated mitochondria (n=4 per condition) from young, young acute ELAM-treated, old, and old acute ELAM-treated skeletal muscle were incubated with a dose of [^3^H]ADP with or without 5 µM carboxyatractyloside (CAT) to inhibit ANT and calculate ANT-specific uptake of [^3^H]ADP. Black bars represent percent of total [^3^H]ADP observed with 5 µM CAT and colored bars represent difference between total [^3^H]ADP and [^3^H]ADP uptake observed with 5 µM CAT, which is specific to flux through the ANT.

**Figure S7. Protein Abundance and Phosphorylation Changes in the ADP/ATP Transport and Synthesis Pathway**

A) Log_2_ fold change of protein abundance of old/young and old with chronic ELAM treatment/old in skeletal muscle and heart for ANT, VDAC, ATP synthase proteins, and creatine kinase proteins for all proteins detected in either tissue.

B) Log_2_ fold change of protein phosphorylation at intensities individual residues of old/young and old with chronic ELAM treatment/old in skeletal muscle and heart for ANT and VDAC proteins for all phosphorylated residues detected in either tissue.

*q<0.05 for each comparison. X values are not detected.

**Supplemental Video 1. Elamipretide Repairs Global Longitudinal Strain (GLS).**

Representative strain analysis of a

A) Young heart,

B) Old heart pre-ELAM treatment, and

C) Old heart post-ELAM treatment. The old pre- and post- videos are from the same mouse.
