## Supplemental Figures for "Elamipretide Improves ADP Sensitivity in Aged Mitochondria by Increasing Uptake through the Adenine Nucleotide Translocator (ANT)"

#### Figure S1. Comparisons of Multivariate Mitochondrial Data

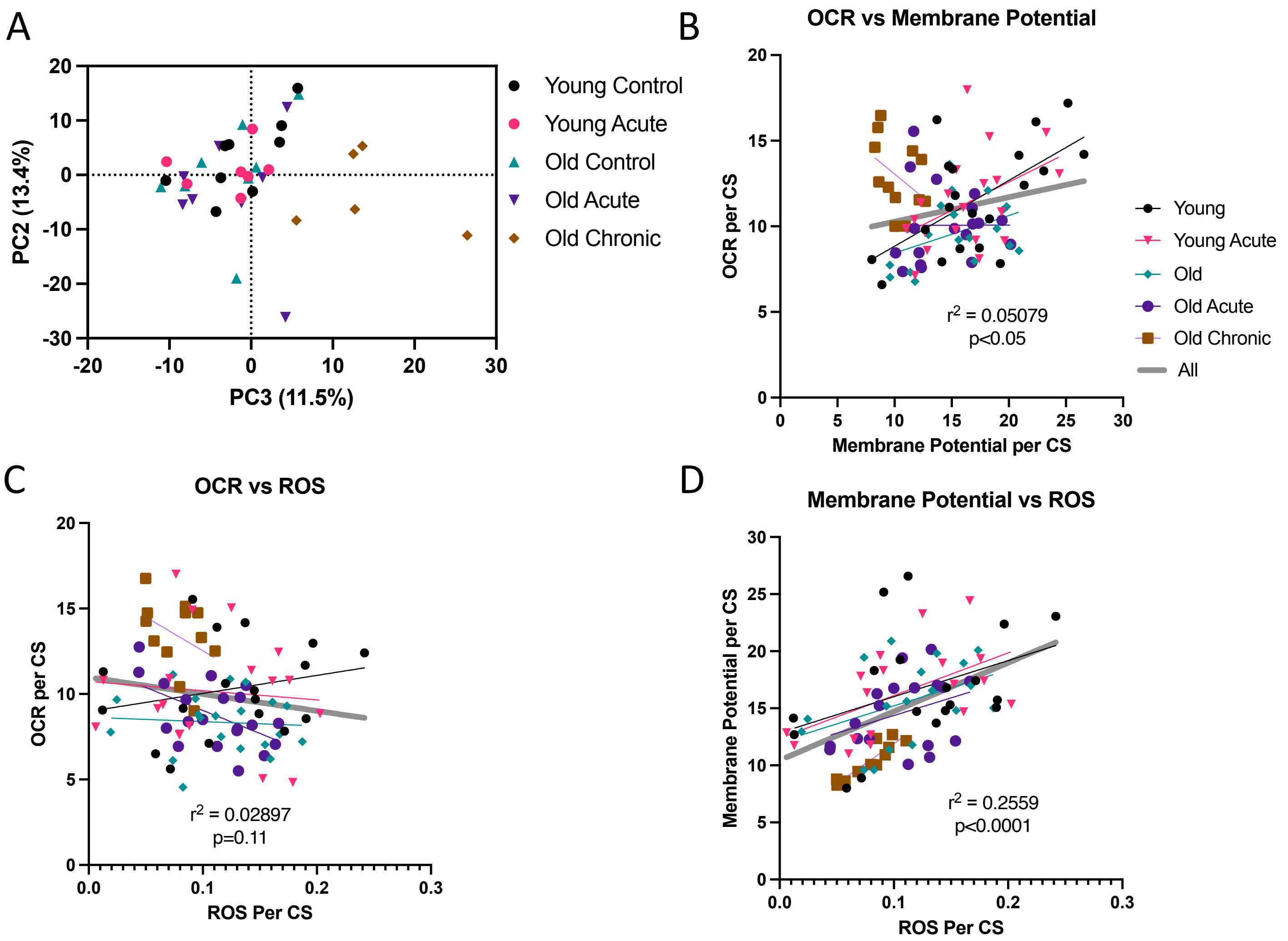

### Figure S2. ADP Kinetic Sensitivity is Not Changed but OXPHOS Capacity is Reduced with Age in Isolated Female Muscle Mitochondria.

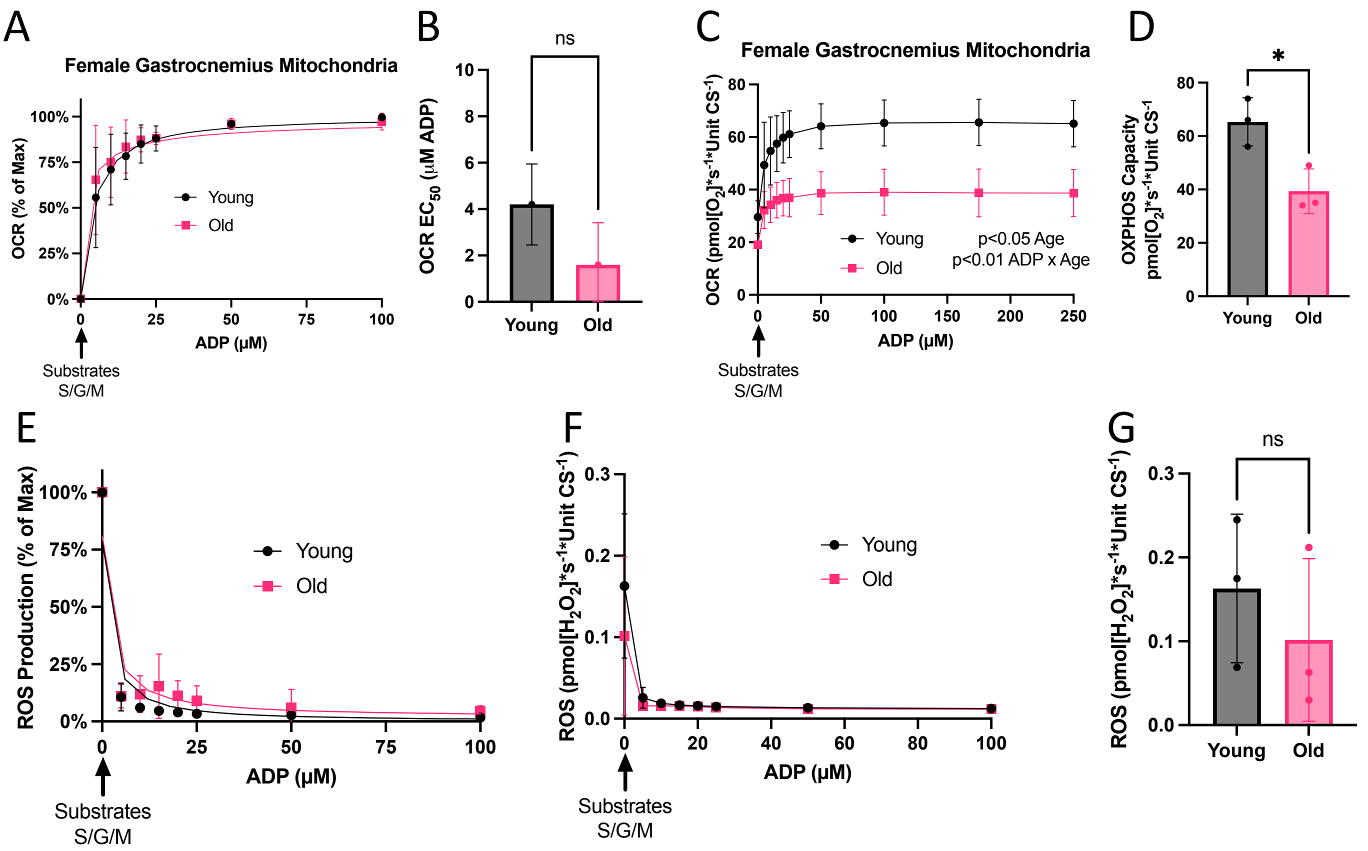

### Figure S3. Acute ELAM Treatment Does Not Affect Young Muscle Mitochondria

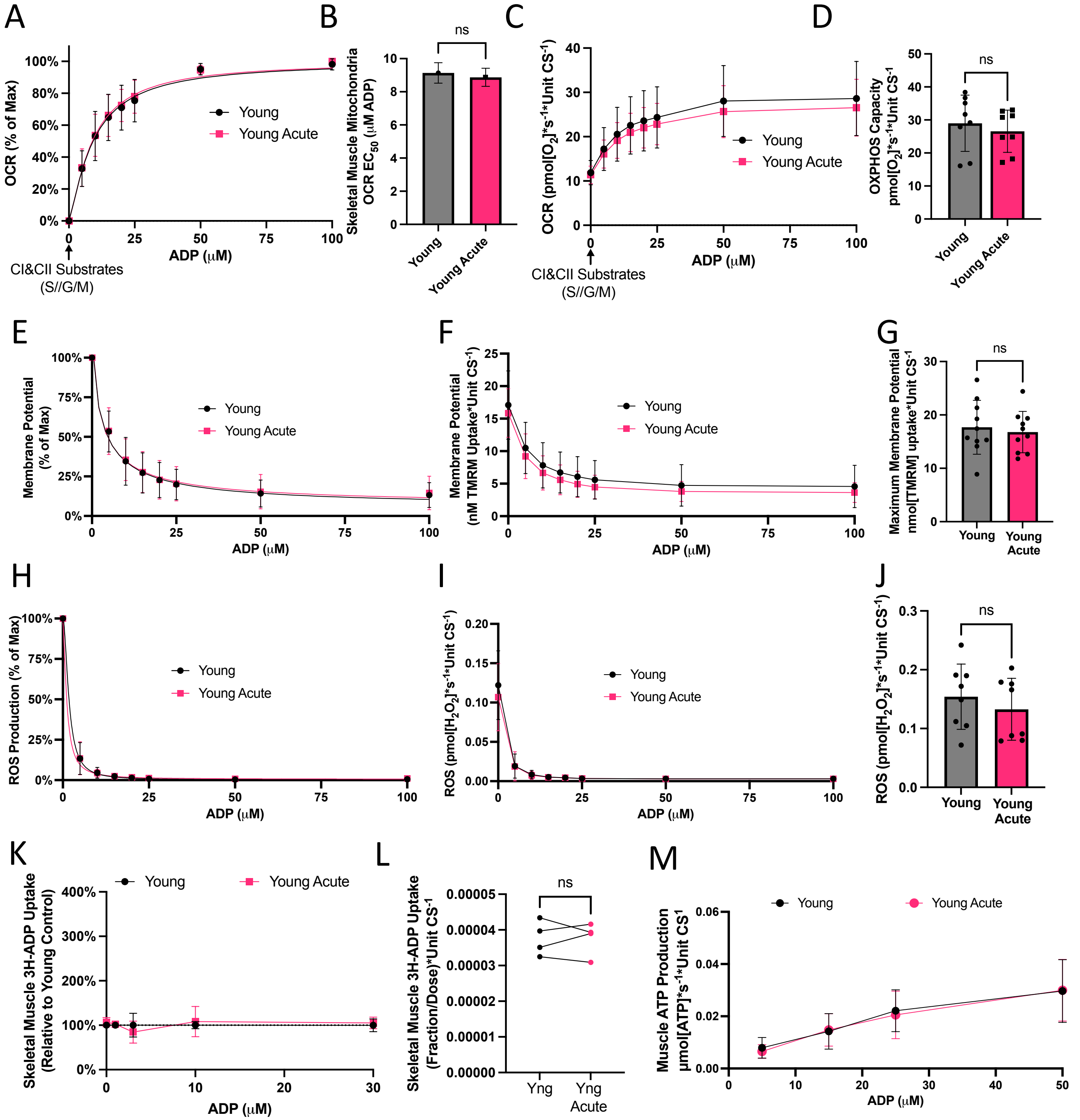

### Figure S4. ELAM Improves ADP Sensitivity for Oxygen Consumption Rate in Aged Isolated Mitochondria Across all Buffer Conditions.

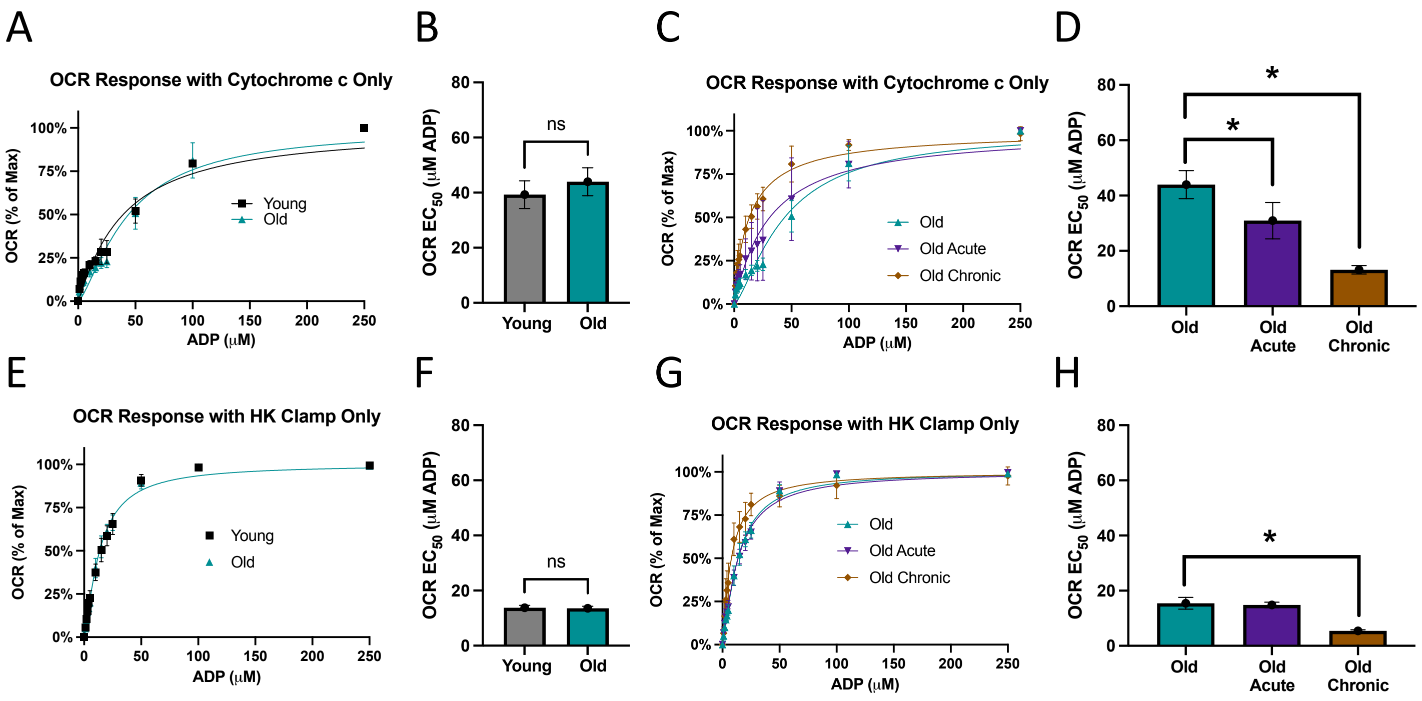

### Figure S5. ELAM Effects on ADP Sensitivity are Replicated in Aged Muscle Fibers

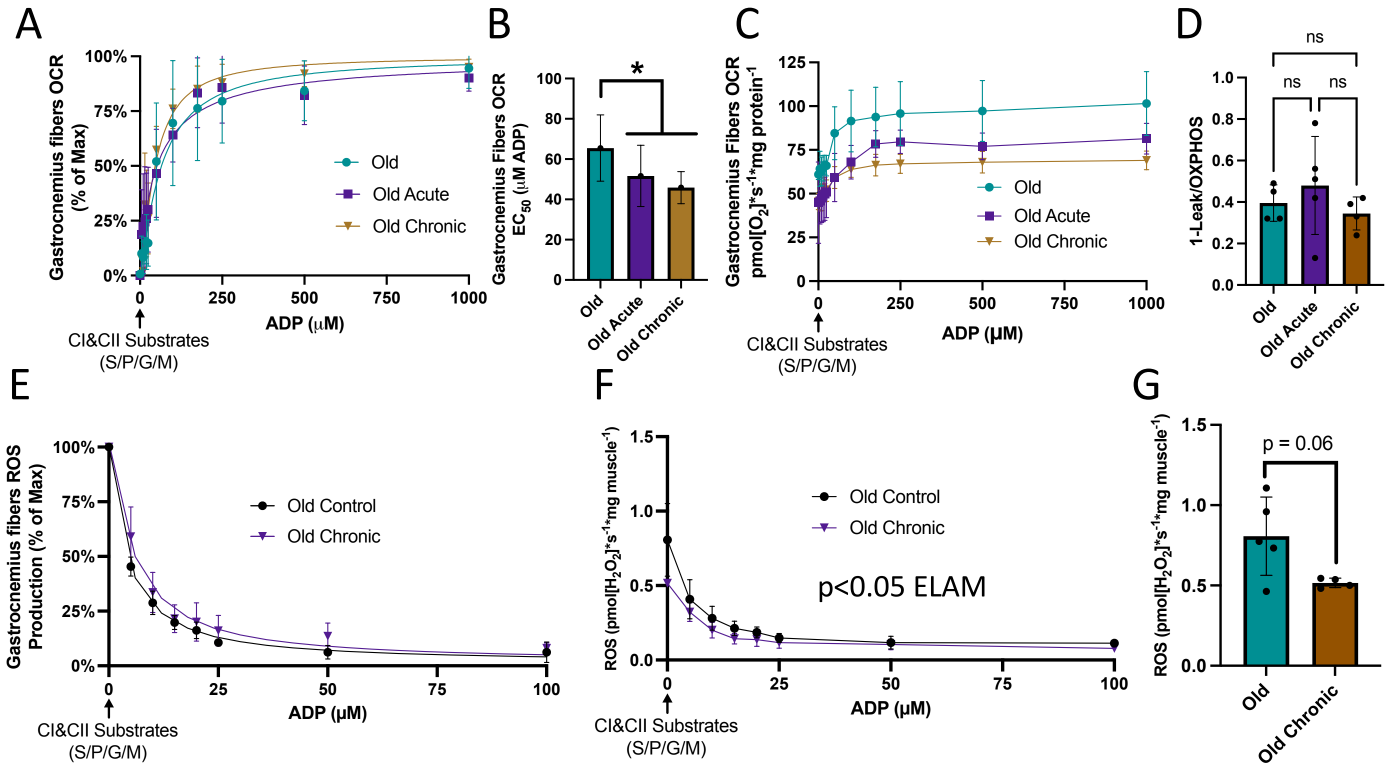

**Figure S6. Proportion of Total [$^3$H]ADP Uptake with and without Carboxyatractyloside (CAT).**

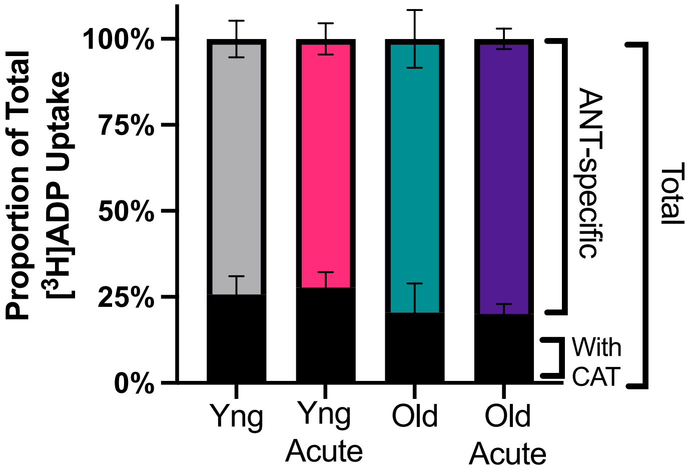

**Figure S7. Protein Abundance and Phosphorylation in the ADP/ATP Transport and Synthesis Pathway**

A

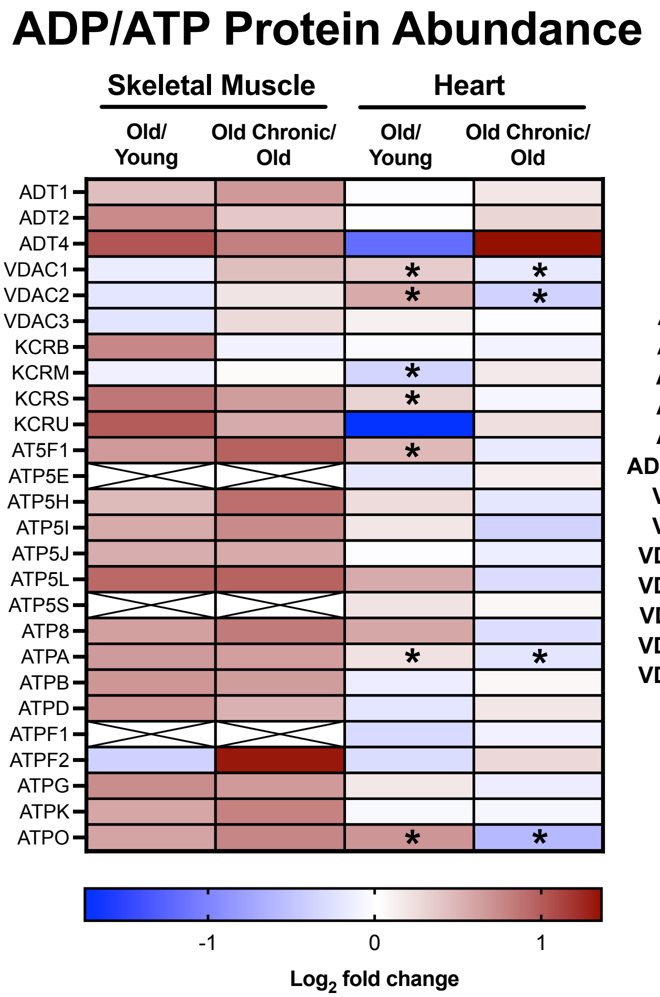


B

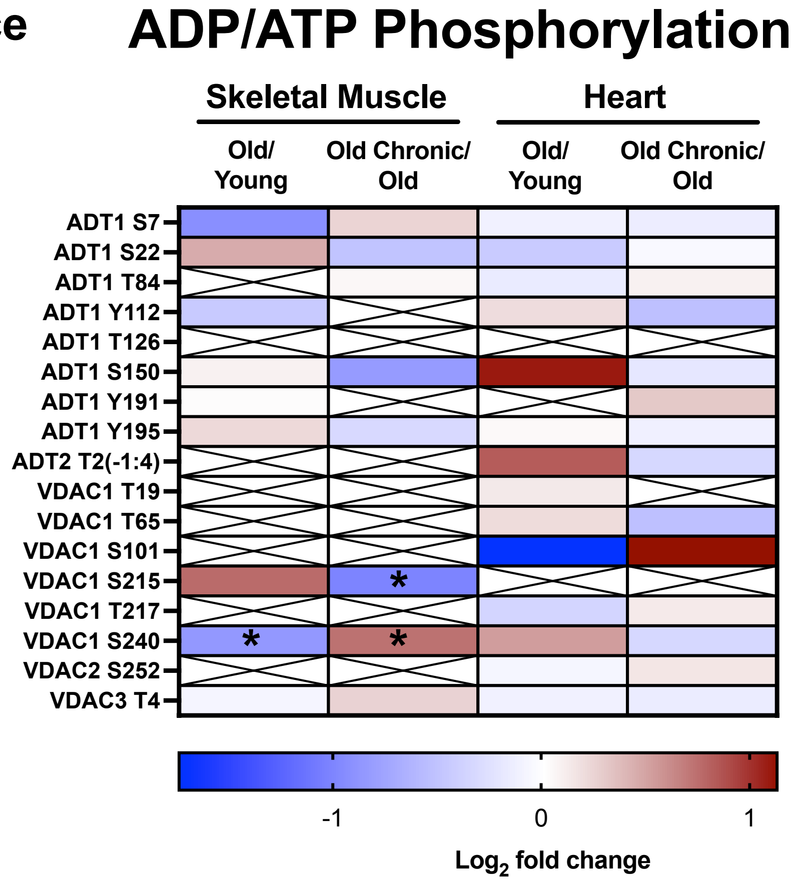
